# CycleVI: Isolating cell cycle variation with an interpretable deep generative model

**DOI:** 10.1101/2025.11.04.686009

**Authors:** Pia Mozdzanowski, Marcel Tarbier, Gustavo S. Jeuken

## Abstract

**Motivation:** Cell cycle progression is a dominant source of variation in single-cell RNA sequencing (scRNA-seq) data, often obscuring other transcriptional signals of interest. Several methods have been developed to infer continuous cell-cycle phase from transcriptomic data, but their estimates tend to be unstable when proliferation is intertwined with other biological processes or technical sources of heterogeneity.

**Results:** We present CycleVI, a deep generative model that disentangles cell cycle–driven variation from other signals in scRNA-seq data using a partitioned latent representation with a dedicated circular subspace. CycleVI accurately infers a continuous cell cycle phase, validated against orthogonal protein-level measurements, and yields a residual latent space free of cell cycle artifacts. This disentangled representation helps resolve biological processes intertwined with the cell cycle, clarifying hematopoietic differentiation and preserving drug-response signals better than standard cell cycle regression. By isolating cell cycle–related variation rather than removing it, CycleVI provides a principled framework for analyzing cellular heterogeneity in proliferating systems.

**Availability and Implementation:** CycleVI is available at www.github.com/jeuken/CycleVI.

## 1 Introduction

Single-cell RNA sequencing (scRNA-seq) has revolutionized the study of complex biological systems by enabling the profiling of transcriptomes from large numbers of individual cells. This technology provides an unprecedented view of cellular heterogeneity. However, this increased resolution of scRNA-seq data is accompanied by substantial methodological challenges [1, 2]. Among these, identifying and disentangling key sources of biological variation from inherently noisy data remains a central challenge [3, 4, 5].

One of the most significant sources of biological variation is the cell cycle, a fundamental process essential for tissue homeostasis and development [6, 7], and whose dysregulation is a hallmark of cancer [8]. Progression through the cell cycle is driven by large, coordinated waves of RNA synthesis and turnover [9, 10], which often constitute the dominant transcriptional signal in scRNA-seq datasets from proliferating cell populations [7]. This high-amplitude variation can obscure more subtle, yet biologically informative, signals related to cell identity, differentiation state, or responses to external perturbations. Consequently, standard bioinformatics workflows that rely on dimensionality reduction techniques to capture the principal axes of variation are prone to organizing cells primarily along the cell cycle axis [11, 12]. If unaccounted for, the cell cycle effect can confound clustering of cell types, distort trajectory inference, and lead to incorrect identification of differentially expressed genes, thereby compromising the fundamental conclusions of a study [13, 14, 15].

Initial computational strategies to mitigate this issue have centered on assigning cells to discrete cell cycle phases based on the expression of canonical marker gene sets, a method popularized by toolkits like Seurat [12] and Cyclone [16]. Although widely adopted, this strategy discretizes an inherently continuous and circular process.

More recent methods aim to infer a continuous cell-cycle trajectory by estimating a pseudotime or phase angle for each cell. DeepCycle [17] and VeloCycle [18] achieve high-resolution reconstructions but critically depend on RNA velocity [19], requiring access to spliced and unspliced transcript counts that are not always available in preprocessed public datasets. Several approaches, including Cyclum [20], Tricycle [21], Revelio [22], and Peco [23], infer a continuous cell-cycle position directly from static scRNA-seq expression profiles. A more comprehensive review of cell-cycle phase identification methods, including a benchmark of selected methods against discrete phase labels, is provided by Guo and Chen [24]. Despite the growing number of available methods, existing methods are primarily designed to estimate cell-cycle position or remove cell-cycle-associated variation, rather than to disentangle periodic cell-cycle expression from other strong, overlapping sources of transcriptional variation under noisy conditions. This leaves a methodological gap for noise-aware approaches that can infer continuous cell-cycle state from static expression data while explicitly separating cell-cycle-driven variation from other biological programs.

Deep generative models, particularly variational autoencoders (VAEs), offer a principled framework for modeling the complex, high-dimensional probability distributions of scRNA-seq data. One important step in this direction was made by the scVI method [25, 26], which demonstrated that VAEs can effectively model technical variability such as gene dropouts and batch effects while learning biologically meaningful latent representations. Importantly, this modeling paradigm can be adapted to move beyond removing confounding variation toward disentangling multiple, overlapping biological signals into a modular and analyzable representation.

Here, we present CycleVI, an interpretable deep generative model that disentangles cell cycle–driven transcriptional variation from other sources of variation in static scRNA-seq data. CycleVI is built on a VAE architecture with a partitioned latent space, comprising a circular subspace that encodes continuous cell cycle phase and a residual subspace capturing remaining variation. Gene expression is reconstructed using a dual-decoder design with a strong inductive bias: a gene-specific Fourier series decoder models periodic, cell cycle–dependent expression, while a neural network decoder models non-cycling variation. An adversarial classifier further enforces separation by discouraging the residual latent space from encoding cell cycle information.

## 2 Methods

### 2.1 Model architecture

Below, we first describe the architecture of CycleVI and then discuss three key modeling choices in detail: (1) how the cell cycle is represented in the latent space, (2) its influence on gene expression, and (3) strategies for addressing the numerical challenges of the architecture.

The architecture of CycleVI is illustrated in Figure 1. CycleVI uses a neural network encoder to map the high-dimensional transcriptome *x*_*n*_ of each cell *n* to the parameters *µ*_*n*_ ∈ ℝ^*D*^ and 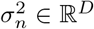 of a multivariate Gaussian distribution over a *D*-dimensional latent space. A latent vector is then sampled as

**Figure 1:**
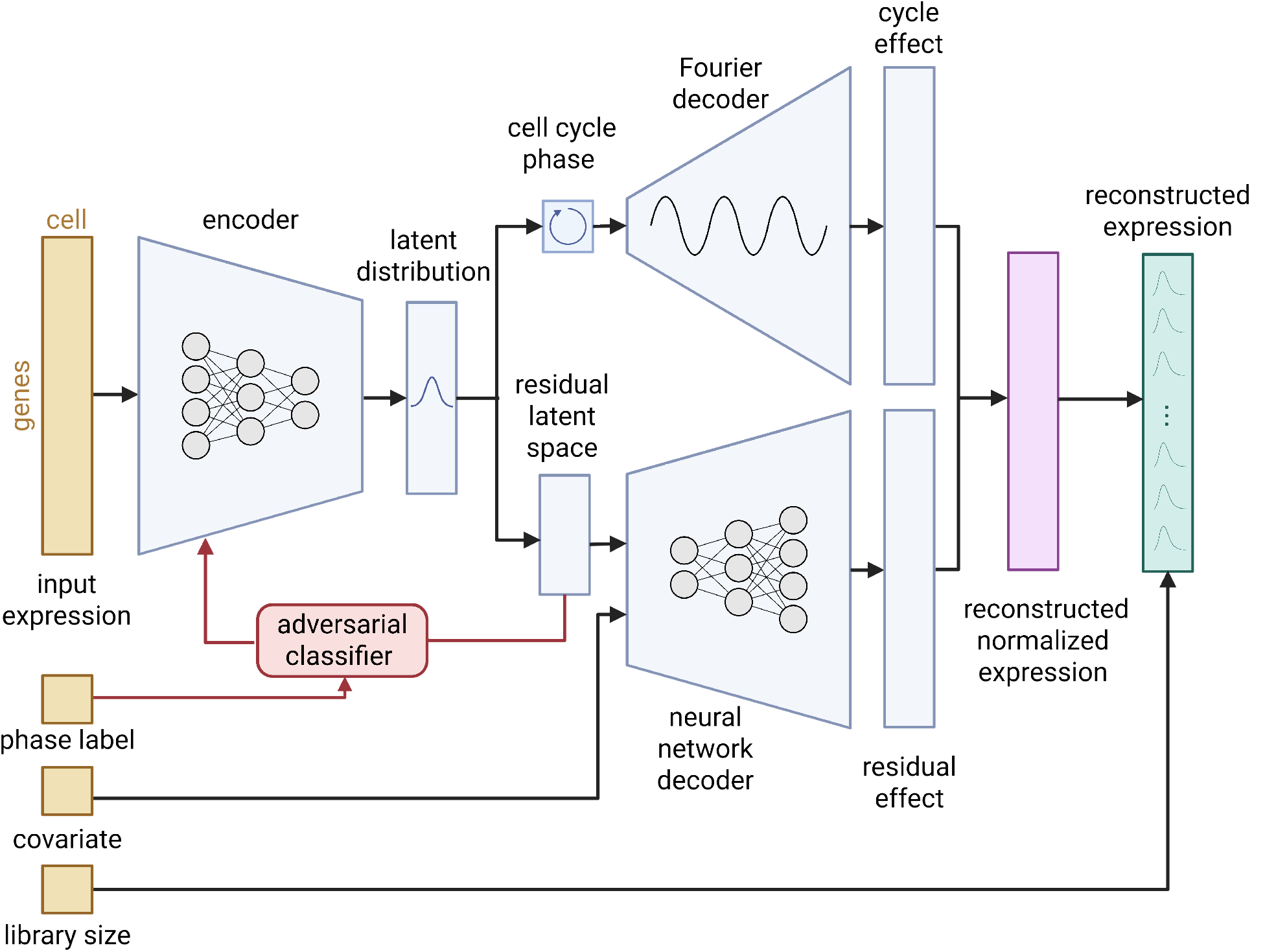
CycleVI architecture. A neural network encoder first maps normalized gene expression data to a low-dimensional multivariate Gaussian distribution, from which a latent representation is sampled. This latent space is then partitioned into a two-dimensional subspace, where the angular coordinate encodes the continuous cell cycle phase, and a residual subspace that captures other sources of variation. To decode the angle of the circular component, a gene-specific Fourier decoder is used to model periodic, cell-cycle-driven expression patterns. In parallel, the residual component, together with observed covariates, is decoded through a neural network to account for the remaining variation. The outputs of both decoders are subsequently combined and passed to a softmax function to obtain the reconstructed normalized expression. Following this, the normalized expression is scaled by the observed library size to yield expression rates. These rates then parameterize a gene-specific negative binomial distribution that models true transcript counts. Finally, an adversarial classifier, trained on categorical cell cycle phase labels (e.g., obtained from Seurat), is applied to the residual latent space to discourage it from encoding cell cycle phase information during training.

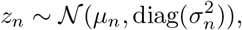

and passed to the generative model. For each gene *g*, the model reconstructs expression in cell *n* through the following probabilistic process:

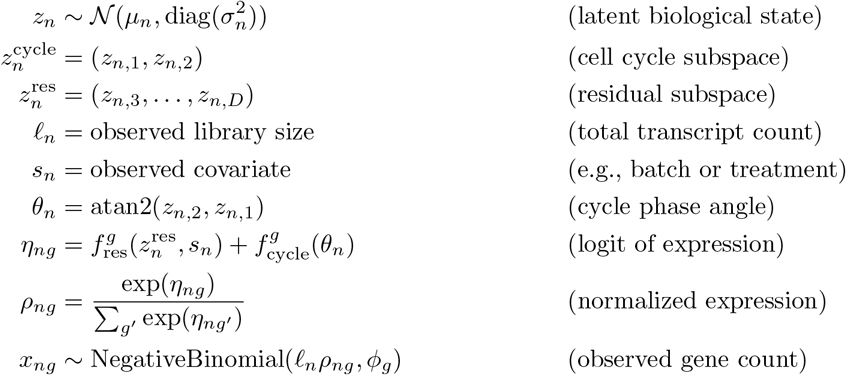

The sampled latent vector *z*_*n*_ ∈ ℝ^*D*^ is partitioned into two subspaces: a two-dimensional component 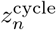, which encodes the position of the cell in the cell cycle, and a complementary subspace 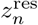 that captures additional sources of biological variation. We use a total latent dimensionality of *D* = 10, with two dimensions allocated to the cell-cycle subspace and the remaining eight forming the residual subspace, following the default convention in scVI [25]. The position in the cycle-specific subspace defines an angular coordinate *θ*_*n*_ relative to the origin, which provides a continuous and circular representation of the cell cycle phase.

The model reconstructs gene expression using two decoder components. The first, a neural network decoder *f*_res_, takes the residual latent representation 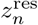 along with optional covariates *s*_*n*_ (e.g., batch or treatment) as input and outputs a gene-specific baseline signal. The second decoder captures periodic modulation as a function of *θ*_*n*_, using a separate truncated Fourier series for each gene, which we explain in more detail in Section 2.1.2. These two components are summed to produce the logit for each gene, which is then passed through a softmax function to obtain the normalized expression proportions *ρ*_*n*_.

The expected expression count for gene *g* in cell *n* is given by the product of the cell’s total transcript count *𝓁*_*n*_ and the normalized expression *ρ*_*ng*_. Observed gene counts are drawn from a negative binomial distribution with mean *𝓁*_*n*_*ρ*_*ng*_ and a gene-specific dispersion parameter *ϕ*_*g*_. A plate diagram of this generative model can be seen in Supplementary Figure S1.

#### 2.1.1 Cell cycle phase model

We represent the cell cycle phase as an angle to account for the periodic and continuous nature of the cell cycle. However, standard variational inference encourages latent vectors to remain close to the origin due to the loss function’s KL divergence term with the standard normal prior. For cells with low radial magnitude, the phase angle *θ*_*n*_ becomes unstable, as small changes to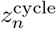 lead to large angular shifts.

To address this, we add a radius penalty that encourages larger norms of 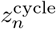, stabilizing the phase representation, and improving interpretability:

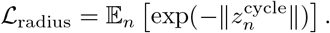

The weight of this term is increased over training, making it more influential as optimization progresses, and thus still giving the model enough autonomy in its initial training steps.

#### 2.1.2 Cell cycle expression model

Once the cell cycle phase is encoded as an angle *θ*, we must model how this cyclic phase modulates gene expression. Due to the periodic nature of the cell cycle, the mapping from phase to expression should also be periodic. While simple sinusoidal functions, such as a single cosine term, offer a natural starting point, they are overly restrictive, capturing only a single, symmetric peak and trough per cycle. This fails to reflect the diversity of periodic expression patterns observed across cell cycle-regulated genes.

To overcome this limitation, we model phase-dependent expression using a truncated Fourier series:

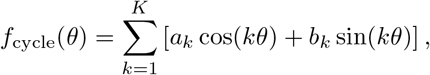

The number of harmonics, K, was set to 3. This value was selected based on a cross-dataset generalization experiment (Supplementary Section 2), in which we trained CycleVI on the hematopoiesis dataset (Section 3.7) with *K* ∈ 1, …, 5 and evaluated how well the learned gene-specific Fourier coefficients transferred to the FUCCI dataset. Performance increased from *K* = 1 to *K* = 3, plateaued at *K* = 3 and *K* = 4, and dropped sharply at *K* = 5, consistent with a bias-variance tradeoff. We selected *K* = 3 as the most parsimonious choice within the plateau region, with 6 coefficients per gene compared to 8 for *K* = 4, and sitting further from the overfitting regime indicated by *K* = 5.

The coefficients *a*_*k*_, *b*_*k*_ are learned per gene. This flexible formulation accommodates a wide range of periodic profiles, including multiple peaks, asymmetries, and phase shifts, but crucially models only one period, thus capturing only the dominant cycling period of the cell. This leads to interpretability and a strong inductive bias toward cyclical structure.

#### 2.1.3 Overcoming optimization challenges with informed initialization

While the approach described above seems intuitive, it presents a great numerical challenge, as the model must simultaneously infer both the cell’s phase *θ* and the function that connects it to the gene expression. This creates a circular dependency: if the latent space does not already organize cells by phase, the decoder receives no coherent signal to learn cyclic modulation; conversely, without a working decoder, the latent space has no incentive to align cells meaningfully along a cycle. This mutual dependency makes the optimization extremely nonconvex, and the training very prone to convergence to poor local optima.

A common issue we faced was that the encoder would collapse the inferred angles to a narrow range so that the decoder could capture non-cyclic variation by using only parts of the periodic function, thereby undermining the intended structure of the latent space.

We can mitigate this problem by an informed initialization of our latent space, biasing it towards the global minimum. This can be done by using an approximate first guess for each cell’s position in the cell cycle. Here, we tested using the expression of single marker genes, of a pool of histone genes, and measures of complexity of expression. Ultimately, we choose Seurat [12, 27] scoring of a panel of marker genes as a well-established and robust method for cell-cycle phase classification. We introduce an auxiliary loss term that is present only at the beginning of training, and aligns the inferred phase angle *θ*_*n*_ with a reference angle 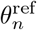, derived from Seurat’s cell cycle scoring method.

Seurat computes two continuous scores per cell, the S and G2/M scores, based on the average expression of curated marker genes specific to the S and G2/M phases [27]. To convert them into a phase angle, we interpret the scores as Cartesian coordinates and compute:

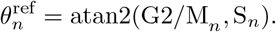

To ensure uniform coverage of the circular domain, we apply a quantile transformation to 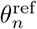 across all cells, spreading the values uniformly over the circle. This mitigates the risk of underfitting the periodic function in sparsely sampled regions of the cell cycle. Importantly, this assumes that the dataset includes cells spanning all phases of the cycle, which may not always hold.

The angle alignment loss is defined as:

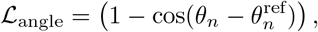

The cosine-based loss penalizes angular deviations and is annealed over time, ensuring that strong supervision is imposed early in training and quickly relaxed, allowing the model to refine the phase assignment. The angle-alignment loss uses a fourth-power annealing schedule (Section 2.2) so that strong supervision is concentrated in the earliest training epochs, where the latent space is most prone to the collapse mode described above, and is rapidly relaxed once the decoder has learned a coherent periodic representation. Continued strong alignment beyond this point would over-constrain the model and prevent it from refining the phase assignment based on the data.

#### 2.1.4 Adversarial separation of latent subspaces

Another challenge of this proposed architecture is the flexibility of the decoder. Because of the universal characteristic of the neural network 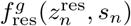, there is nothing to stop it from also capturing the variability attributed to the cell cycle, which would defeat the purpose of the architecture.

To enforce disentanglement between the cell cycle and residual latent subspaces, we use an adversarial classifier. To this end, we first label the samples according to an initial guess for their cell cycle phase, and here we use Seurat’s phase assignments (G1, S, G2/M) for consistency. We then train an adversarial classifier to predict this phase label from the residual latent space 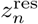, while simultaneously training the encoder to make this prediction difficult. Specifically, the adversarial classifier minimizes cross-entropy with the true label, while the encoder minimizes a softened cross-entropy with a target distribution uniform over the incorrect classes:

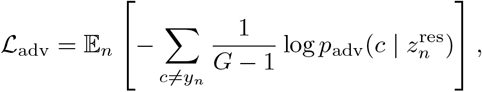

where *y*_*n*_ ∈ {G1, S, G2/M} is the initial label, *G* = 3, and *p*_adv_ is the output of the classifier. This loss encourages phase-related information to be captured in its own subspace, while pushing the residual representation toward phase invariance. The adversarial term is annealed over training and deactivated by the final epoch, allowing the model to fine-tune expression reconstruction without being constrained to enforce strict independence between the cycle and residual subspaces, an assumption that may not hold perfectly due to biological interactions between the cell cycle and other transcriptional programs.

### 2.2 Training

The model is trained using stochastic variational inference to maximize the Evidence Lower Bound (ELBO), augmented with three auxiliary objectives. Optimization is performed using the Adam optimizer with a learning rate of 10^*−*3^, mini-batch size of 128 cells, and a total of *T* = 400 training epochs.

At epoch *t*, the total objective minimized is:

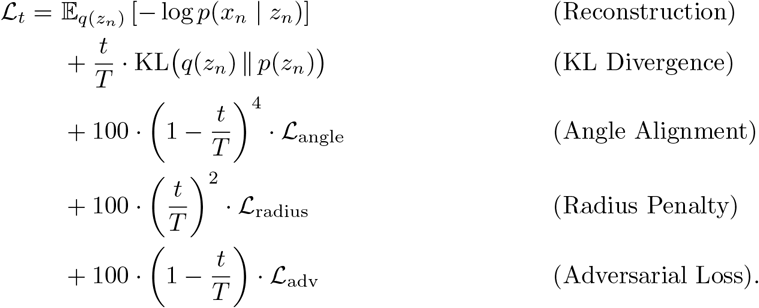

The four auxiliary terms follow different annealing schedules tailored to their roles in optimization. The angle-alignment loss uses a fourth-power decay because it is needed only to break the symmetry of the optimization at the start of training; once the decoder has learned a coherent periodic representation, continued alignment becomes counter-productive. The adversarial loss follows a slower linear decay because phase information can leak into the residual subspace for longer in training, until the cell-cycle decoder has approximately fitted the periodic effect. The radius penalty grows over training (linear schedule), becoming most influential once the angular structure is established and stable angle estimation matters most. The KL term uses linear annealing, the standard schedule for VAEs, gradually enforcing the prior as training progresses.

## 3 Results

To validate and demonstrate the utility of our method, we applied it to six datasets, described in Supplementary Section 1. We first verify that the inferred phase recovers known cell-cycle transcriptional dynamics (Section 3.1). We then benchmark CycleVI against seven established methods using an orthogonal FUCCI protein-level ground truth, where it achieves the highest circular rank correlation on both a controlled and a heterogeneous dataset (Section 3.2). We show that CycleVI distinguishes proliferating from quiescent populations in FFPE tissue (Section 3.3) and cleanly partitions periodic from non-periodic expression in its latent space (Section 3.4). In a chemical perturbation setting, CycleVI removes cell-cycle structure more completely than standard linear regression while better preserving drug-response signals (Section 3.5). Finally, we demonstrate its applicability to spatial transcriptomics (Section 3.6) and show that disentangling the cell cycle clarifies the human hematopoietic differentiation manifold (Section 3.7). Supplementary analyses establish robustness to noisy initialization (Supplementary Section 3), agreement with velocity-based methods on static data (Supplementary Section 4), and improved generalization relative to scVI through the incorporation of biological inductive biases (Supplementary Section 5).

### 3.1 CycleVI inferred progression aligns with known transcriptional dynamics of the cell cycle

As an initial sanity check that the inferred phase angle reflects cell-cycle biology rather than other sources of variation, we examined whether canonical cell-cycle markers organize along the angle in their expected temporal order. Figure 2a displays the expression profiles of selected canonical cell cycle genes along the inferred phase angle (more markers are displayed in Supplementary Figure S2). These markers exhibit clear periodic expression patterns consistent with their established roles in the cell cycle [28]. For example, Cyclin E (CCNE1) peaks at the G1/S transition, promoting DNA replication by activating CDK2 and inducing S phase genes such as PCNA. Cyclin A2 (CCNA2) accumulates throughout S and G2 phases to support replication completion and entry into mitosis. Cyclin B1 (CCNB1) and TOP2A peak at G2/M, facilitating mitotic entry and chromosome condensation.

**Figure 2:**
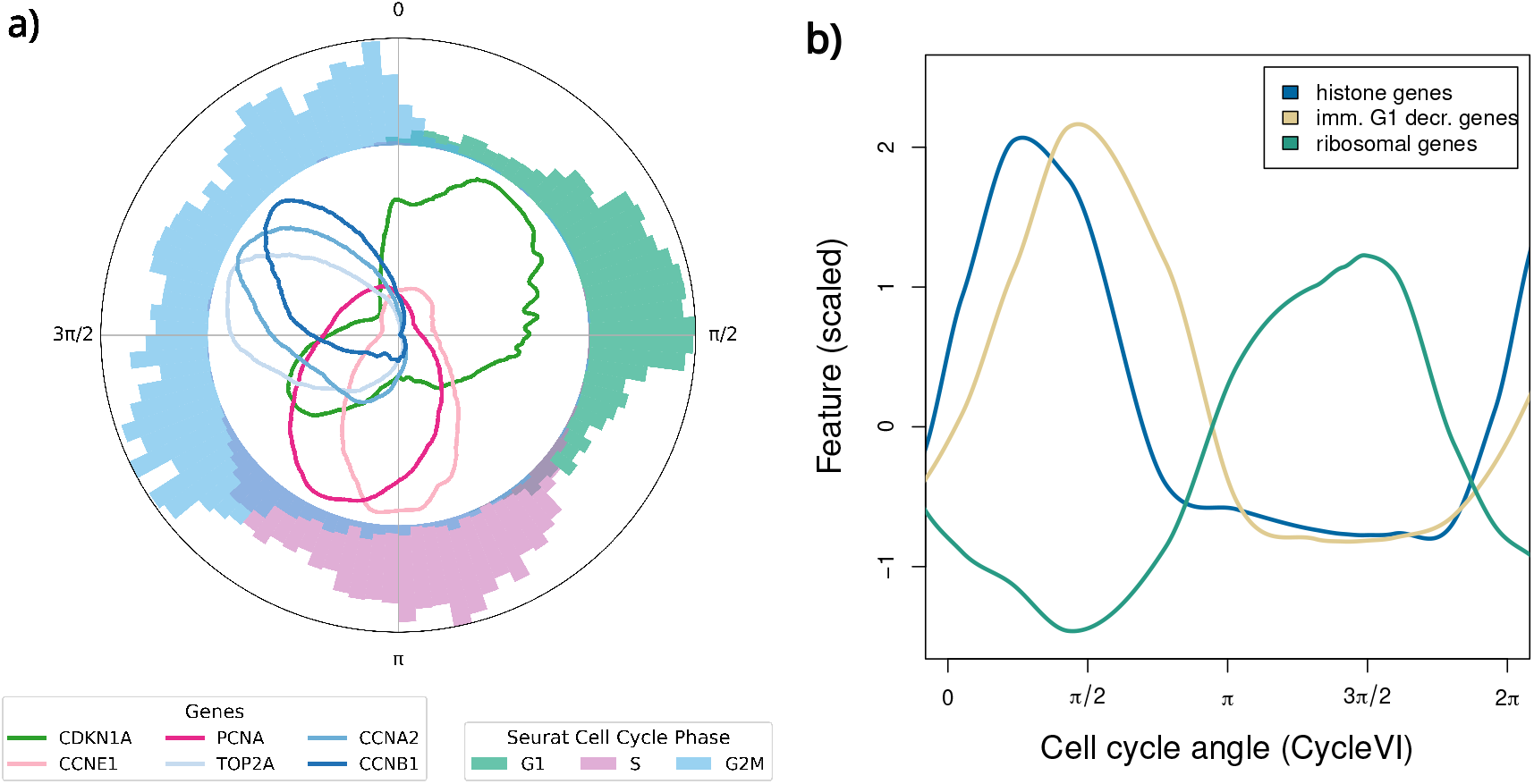
a) Expression of selected marker genes plotted against the phase angle inferred by CycleVI. Each curve represents the Gaussian-smoothed (*σ* = 50), min-max scaled, normalized expression of a gene. The outer histogram shows the distribution of Seurat-assigned phase labels (G1, S, G2/M), indicating how discrete annotations align with the learned continuous trajectory. The alignment of gene expression peaks with known cell cycle transitions, such as CCNE1 and PCNA in S phase or CCNB1 and TOP2A in G2/M, validates the inferred phase ordering. **b) Global gene expression changes and alternative markers**. Scaled LOESS fit for the fraction of histone genes, of genes decreasing in early G1 phase from Krenning et al. [9], and of ribosomal protein genes across estimated cell cycle angle.

p21 (CDKN1A), a cyclin-dependent kinase inhibitor of CDK2, exhibits two distinct expression peaks along the inferred cell cycle trajectory: one during the G1 phase and another in G2. The first peak likely reflects the entry of cells into quiescence (G0), consistent with the role of p21 as a well-established marker of quiescent states. Since CycleVI models the cell cycle as a continuous latent trajectory rather than a series of discrete phases, it does not explicitly separate G0 from G1. Nevertheless, the pronounced CDKN1A expression peak in the G1/G0 region may suggest that a substantial fraction of these cells are quiescent. This interpretation aligns with findings from the developers of DeepCycle, who reported a high prevalence of quiescent cells in this dataset [17]. The second peak in CDKN1A expression during G2 likely corresponds to its established function in enforcing the G2 DNA damage checkpoint.

Figure 2b shows how CycleVI accurately captures the known expression dynamics of major transcriptomic events across the cell cycle. The model correctly identifies the increase of histone gene transcripts during S-phase and their subsequent clearance at S-phase exit, as well as the robust increase of ribosomal protein transcripts required for growth in early G1 phase. Additionally, we investigated a small set of transcripts that have been reported to decrease immediately after metaphase [9]. The clearance of these G2/M-specific transcripts serves to reset the cell for the next cell cycle. CycleVI again captures this decrease which is offset from the decrease of histone genes at the exit from S-phase. Overall, the inferred trajectory recapitulates known cell cycle-dependent expression dynamics, supporting the validity of the learned phase. These results indicate that CycleVI is capable of inferring cells’ approximate position in the cell cycle.

### 3.2 CycleVI phase inference agrees with FUCCI protein-level measurements and outperforms other methods

Validation using marker genes can be self-referential, as it uses transcript levels to validate a cellular state defined by those same transcripts. To avoid this circularity, we benchmarked CycleVI against the Battich et al. dataset [29], which provides a non-transcriptional ground truth. This dataset uses the Fluorescent Ubiquitination-based Cell Cycle Indicator (FUCCI) system, which yields a continuous, protein-level estimate of each cell’s position based on the post-translational regulation of two fluorescent proteins. This orthogonal readout enables an unbiased evaluation of cell cycle inference methods from gene expression alone.

The Battich et al. dataset is comprised of two subsets both featuring RPE1-FUCCI cells but varying with regard to depth, variation and cell cycle distribution. The pulse experiments (*n* = 2, 793 cells) comprise cells labeled with 5-ethynyl-uridine for varying times between 15 minutes and 3 hours. The chase experiments extend this with cells labeled for 22 hours followed by 0 to 6 hours of chase with unlabeled uridine. These different labeling strategies are liable to introduce distinct subtle technical biases, batch effects. For instance, while the pulse dataset shows higher sequencing depth, lower variation and a cell cycle distribution biased towards G1 phase. The full dataset combines both (*n* = 4, 955 cells) and therefore spans a broader range of transcriptional and technical variation, making it a more challenging benchmark. We applied each method separately to the pulse subset and to the full dataset. The data contain both spliced and unspliced transcripts, the expression of which we summed when providing input to CycleVI.

We compared CycleVI against seven published cell cycle inference methods, spanning three methodological families. Revelio [22], Tricycle [21], and Scanpy [30] (which implements Seurat [12] cell cycle scoring) rely on curated panels of marker genes. Cyclum [20], like CycleVI, is an autoencoder with a circular latent variable, but does not incorporate biological inductive biases. VeloCycle [18] and DeepCycle [17] require both spliced and unspliced transcripts and perform RNA velocity analysis prior to phase estimation. Peco [23] is a Bayesian phase estimator based on marker genes. To quantify agreement between inferred angles and FUCCI-derived positions, we used the circular rank correlation coefficient of Mardia and Jupp [31], which is the directional-statistics analogue of Spearman’s rank correlation. This metric is invariant to rotations, reflections, and any monotonic transformation of either angle, and is therefore the appropriate choice when comparing pseudotime-like outputs whose absolute scale is not biologically identifiable.

Figure 3 summarizes the benchmark results. CycleVI achieves the highest circular rank correlation on both datasets, with *ρ*_*c*_ = 0.74 on the pulse subset and *ρ*_*c*_ = 0.62 on the full dataset. On the pulse subset it is narrowly ahead of Revelio (0.73) and Cyclum (0.70). On the full dataset it is substantially ahead of Tricycle (0.53), Revelio (0.52), and Scanpy (0.49). CycleVI is ranked first on both regimes.

**Figure 3:**
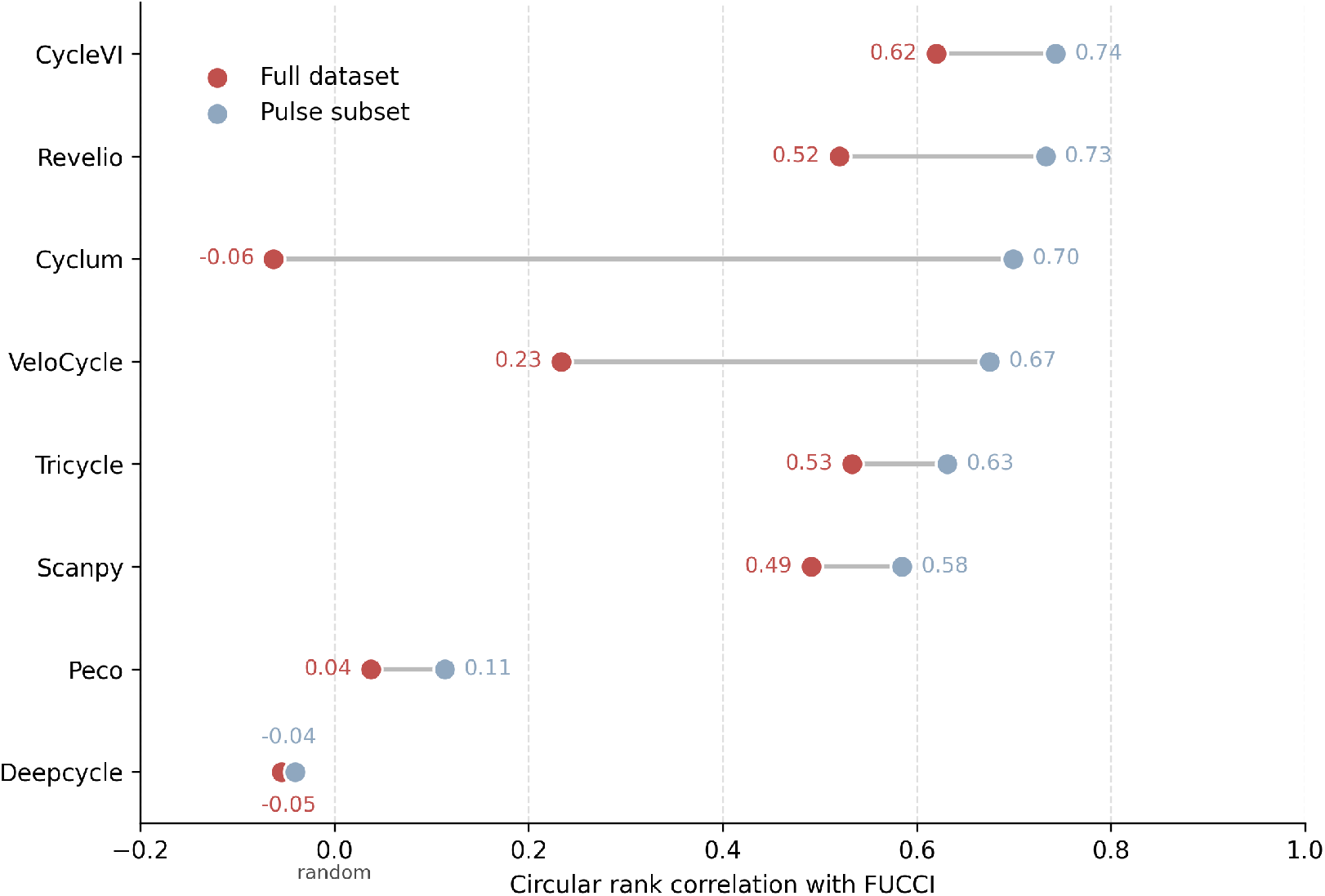
CycleVI outperforms existing cell cycle inference methods against an orthogonal FUCCI ground truth, and is robust to combined experimental heterogeneity. Circular rank correlation between each method’s inferred cell cycle angle and the FUCCI-derived position on the Battich et al. dataset. Each method is shown as a pair of points: the pulse subset (*n* = 2, 793 cells, blue) and the full dataset including the chase experiments (*n* = 4, 955 cells, red). Methods are ordered by performance on the pulse dataset, with the best method at the top. The value of zero corresponds to no association between the inferred phase and the FUCCI ground truth. CycleVI is ranked first on both datasets, and its performance does not degrade substantially when the dataset is expanded to include the chase experiments. Methods that rely on fixed marker-gene panels (Revelio, Scanpy, Tricycle) show moderate degradation, consistent with their inability to absorb batch-level variation. VeloCycle, which assumes a single velocity field shared across all cells, drops sharply. Cyclum, an autoencoder-based method architecturally similar to CycleVI but lacking its biological inductive biases, collapses to near-random performance.

Figure 3 also shows that methods differ substantially in how they behave when moving from the pulse subset to the full dataset. CycleVI’s circular rank correlation decreases only modestly (0.74 to 0.62), whereas several competing methods degrade substantially or collapse entirely. Revelio drops from 0.73 to 0.52, VeloCycle from 0.67 to 0.23, and Cyclum from 0.70 to -0.06. Peco and DeepCycle remain near the null value of zero across both regimes. Among the eight methods tested, CycleVI is the one that retains most informative performance when the dataset is expanded to include the chase experiments. Supplementary Figures 6 and 7 show the results in both experiments for each method.

These differences reflect how each method’s assumptions interact with the heterogeneity introduced by combining the pulse and chase experiments. Marker-based methods (Revelio, Scanpy, Tricycle) assume that a fixed panel of marker genes provides a reliable readout of phase across the dataset. They have no mechanism to absorb batch-level variation, so technical differences between the two subsets enter directly into the phase assignment. Their degradation is moderate rather than catastrophic, since the marker signal itself remains a strong constraint on the inference. Velocity-based methods (DeepCycle, VeloCycle) assume consistent spliced to unspliced dynamics across all cells. VeloCycle in particular fits a manifold-constrained RNA velocity model with a single velocity field shared across all cells [32]. Combining the pulse and chase experiments introduces systematic heterogeneity in the spliced to unspliced signal between the two subsets, which is incompatible with this shared-field assumption, and the resulting distortion in the velocity estimate propagates to the inferred phase. VeloCycle’s drop from 0.67 to 0.23 is the sharpest among methods that do not fully collapse.

Cyclum is the most instructive comparison. Like CycleVI, Cyclum is an autoencoder with a circular latent variable. Unlike CycleVI, it does not incorporate biological inductive biases, and relies on the model’s flexibility to discover the cell cycle from data. In principle, such a model should accommodate heterogeneity by modeling it in additional latent dimensions. In practice, without a principled partitioning of the latent space, the encoder has no incentive to route cycle and non-cycle variation into separate subspaces, and can fit spurious structure when the data is heterogeneous. Cyclum’s collapse from 0.70 to -0.06 indicates that the autoencoder framework alone is not sufficient.

CycleVI incorporates several design choices that address these failure modes. The partitioned latent space constrains cell cycle variation to a dedicated circular subspace, while residual heterogeneity is absorbed by a separate subspace. Observed covariates are passed directly to the residual decoder, providing an explicit mechanism to absorb batch or treatment effects. These choices give CycleVI a principled way to separate cycle signal from technical and biological heterogeneity, which is not a feature of the competing methods benchmarked here.

A reasonable concern with CycleVI is that its performance might be bounded by that of Seurat itself, since the Seurat-based angle 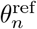 is used to initialize training through the angle-alignment loss. The benchmark results argue against this interpretation. Scanpy, which implements the same Seurat scoring that CycleVI uses for initialization, achieves circular rank correlations of 0.58 and 0.49 on the pulse and full datasets, respectively. CycleVI achieves 0.74 and 0.62, corresponding to relative improvements of 28% and 27% over its own starting point, thus clearly demonstrating its ability to learn cycling gene expression beyond its initialization.

We also note that CycleVI is designed for moderately sized to large scRNA-seq datasets, its variational and adversarial components benefit from sample sizes in the thousands, and on very small datasets (tens to hundreds of cells) simpler marker-based methods such as Seurat scoring may be more appropriate.

### 3.3 CycleVI accurately segregates proliferating tumor cells from quiescent immune populations

We further investigate if CycleVI can perform well when a large portion of non-cycling cells are present. Figure 4 shows that it can distinguish cycling and non-cycling cell populations even in challenging Formalin-Fixed Paraffin-Embedded (FFPE) samples, taken from Janesick *et al* [33]. In a complex FFPE breast cancer biopsy, which contains an intricate mixture of cell types with diverse proliferative activities, CycleVI demonstrates its ability to accurately parse these distinct populations. The model successfully assigns non-cycling, quiescent immune lineages to the G1/G0 phase. This is visually confirmed by a sharp, concentrated peak in their cell cycle angle distribution, which reflects their non-proliferative state. In contrast, the mitotically active tumor cell populations exhibit a pan-phasic and asynchronous distribution, with cells spread broadly across the entire cell cycle angle profile. This pattern is a direct reflection of the uncontrolled and continuous proliferation.

**Figure 4:**
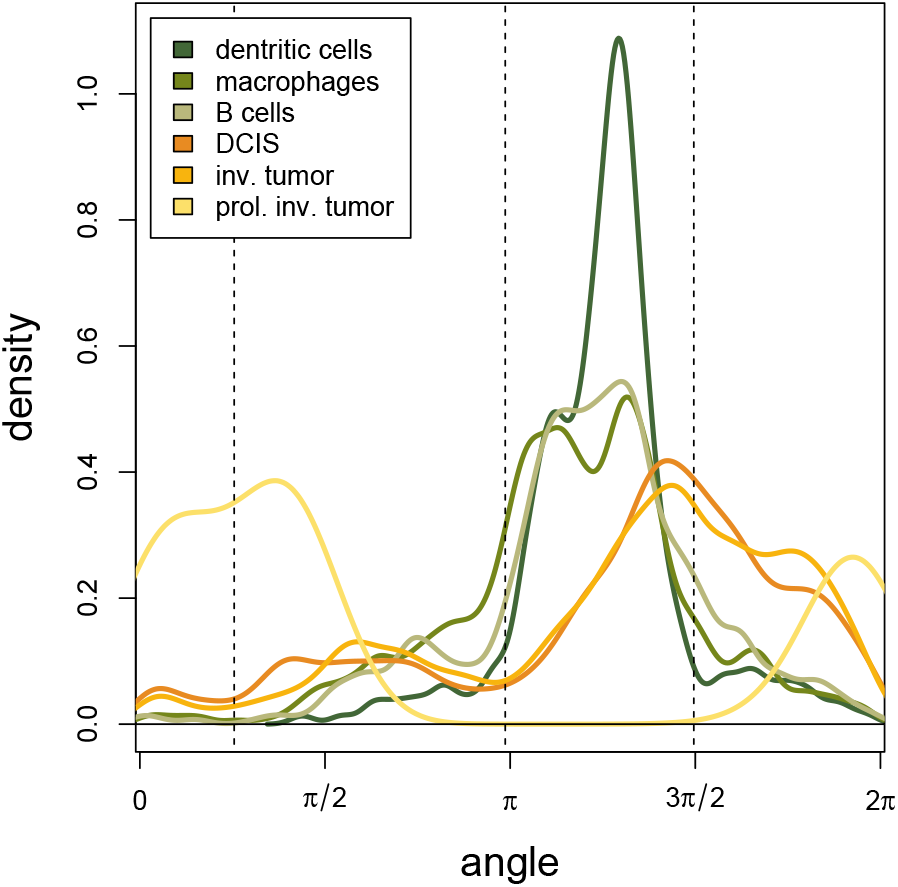
Cycling and non-cycling cell populations in FFPE tissues (breast cancer biopsy). Distribution of cells of various cycling (tumor cells: DCIS, invasive tumor, and proliferative invasive tumor) and non-cycling (immune cells: dendritic cells, macrophages, and B cells) cell types across estimated cell cycle angles.

Beyond this broad separation, CycleVI’s high-resolution continuous phase inference captures subtle but significant heterogeneity even within the malignant cells themselves. It distinguishes between “invasive tumor” cells and a more aggressive “proliferative invasive tumor” subpopulation. This latter group shows a markedly depleted density of cells in the G1 phase region of the cycle, a quantitative observation that suggests an accelerated G1/S transition.

### 3.4 CycleVI successfully isolates cell cycle variation into a dedicated latent subspace

To verify that CycleVI’s architecture achieves the intended decomposition, with cell-cycle variation captured in the dedicated subspace and absent from the residual subspace, we examined the latent representations learned by the model (Figure 5). The goal of this analysis is to validate the model’s internal disentanglement rather than to benchmark cycle removal against alternative methods; the latter comparison is the subject of Section 3.5.

**Figure 5:**
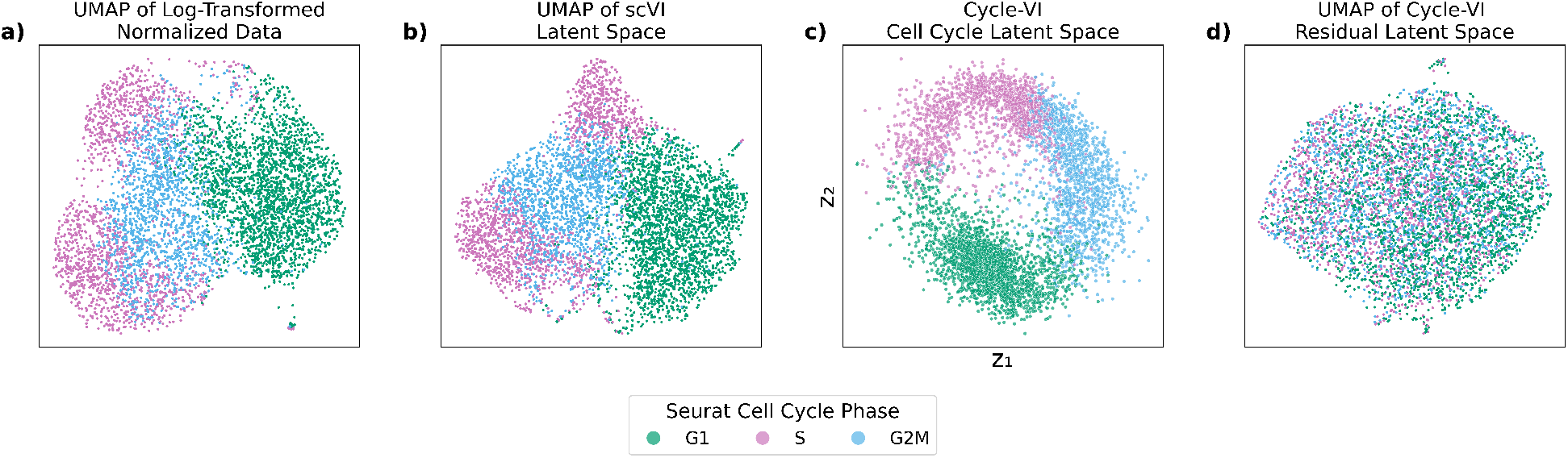
CycleVI isolates cell cycle variation into a dedicated latent subspace. **a)** UMAP of normalized and log-transformed expression data from human fetal lung fibroblasts reveals clustering by Seurat cell cycle phase. **b)** UMAP of the 10-dimensional scVI latent space confirms that the phase is a dominant source of variation. **c)** The dedicated 2D cell cycle subspace in CycleVI captures phase structure explicitly. **d)** UMAP of the 8-dimensional CycleVI residual latent space shows no apparent phase separation, indicating successful disentanglement.

Figure 5a shows that in a UMAP of normalized gene expression values in the fetal lung fibroblast dataset, cells are strongly separated by cell cycle phase. This confirms that phase is a dominant axis of variation in the dataset. Similarly, in the scVI latent space (Figure 5b), cells clearly cluster by cell cycle.

CycleVI learns a dedicated two-dimensional subspace for cell cycle progression (Figure 5c), which cleanly separates cycle phases. The remaining latent dimensions (Figure 5d) no longer exhibit phase structure, suggesting that the model has successfully separated phase-related variation from other transcriptional programs.

To further validate the disentanglement of cell cycle-specific expression, we examined how CycleVI partitions gene expression across its two decoders: one dedicated to modeling cell cycle-driven dynamics (the Fourier decoder), and one modeling the remaining variation (the neural network decoder). As shown in Figure 6, the Fourier decoder output for cell cycle marker genes exhibits a strong periodic structure along the inferred phase, and agrees with the expected expression patterns of these markers (see Section 3.1). In contrast, the non-cycle decoder produces signals that show little to no dependency on the cell cycle angle, confirming that phase-related dynamics are not redundantly encoded by both decoders. The top row illustrates the observed and predicted normalized gene expression, smoothed across the inferred phase angle. The close correspondence between them demonstrates that CycleVI accurately captures the cyclical transcriptional programs.

**Figure 6:**
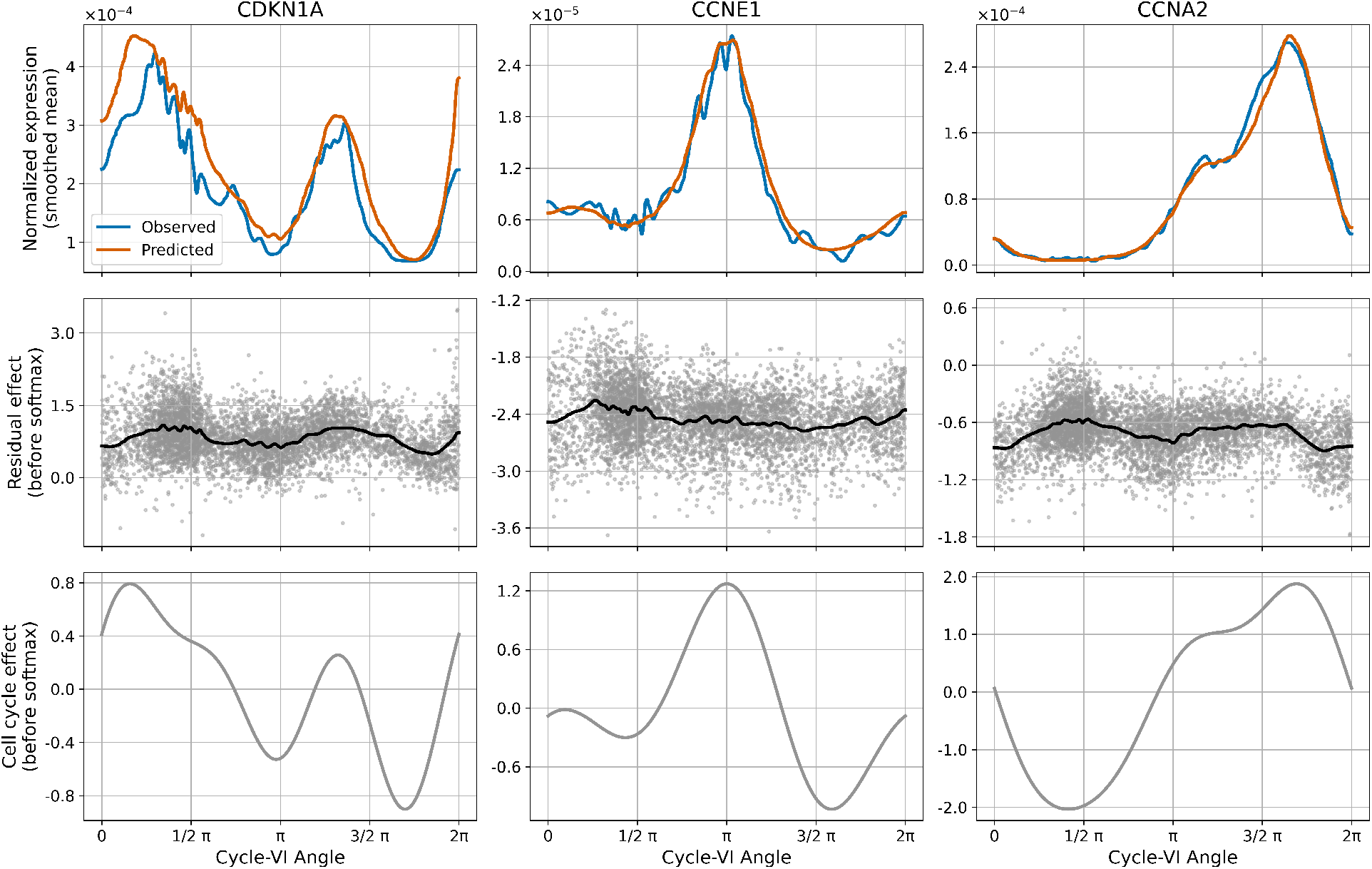
The cell cycle-dependent expression is modeled by the cycle-specific Fourier decoder. Each column represents a selected gene, with the top row showing the normalized observed and predicted expression across the inferred cell cycle angle with Gaussian smoothing (*σ* = 50), the middle row showing the non-cell cycle neural network output before softmax, with a line showing the mean, and the bottom row showing the cell cycle-specific Fourier effect.

### 3.5 CycleVI outperforms linear cycle regression at disentangling drug response from proliferation

A natural question is whether CycleVI’s disentanglement provides meaningful advantages over simpler correction strategies. The standard approach is to compute per-cell S and G2/M scores from curated marker genes and linearly regress them out of the expression matrix, as implemented in Seurat [12]. This procedure can only remove the component of cell cycle variance that is linearly aligned with these two scores. More fundamentally, it is a two-step procedure: phase scores are computed once from a fixed marker panel, and then the gene expression matrix is corrected against those scores in a second, independent step. The phase estimate cannot improve based on what would best explain cycle-driven gene expression in the dataset at hand, and the correction cannot adapt to a better phase estimate. The cell cycle, however, is a circular process whose transcriptional signature is inherently nonlinear: cells progress along a closed trajectory in which the same gene can peak and trough multiple times, and the geometry of this trajectory is not captured by two scalar covariates. Any structure orthogonal to the S and G2/M score directions, but still driven by the cycle, will survive regression and continue to shape the embedding for downstream analysis.

To examine this limitation in a setting where its consequences can be measured, we analyzed a subset of the sci-Plex3 chemical transcriptomics dataset [34], focusing on A549 lung adenocarcinoma cells treated with 9 HDAC inhibitors at four doses (10*nM* to 10*µM*) alongside vehicle controls. HDAC inhibitors induce G1 and G2/M arrest, upregulate the cyclin-dependent kinase inhibitor CDKN1A, and suppress E2F target genes [35], so their transcriptional signature partially overlaps with cycle-regulated programs. This overlap provides a controlled test of whether cycle correction damages or preserves a biological signal of interest.

We compared three representations of the data at matched dimensionality: (i) log-normalized expression followed by principal component analysis (uncorrected); (ii) Seurat cycle score regression followed by PCA (Seurat-regressed); and (iii) CycleVI’s residual latent space. For each, we trained cross-validated predictors to assess cycle removal and biology preservation, using linear (multinomial logistic and linear regression) and nonlinear (k-nearest-neighbor, with *k* = 30) probes. The linear probes assess whether cycle or drug information is linearly recoverable from the embedding, while the nonlinear probes assess whether this information persists in the geometry of the embedding regardless of its linear structure.

Figure 7 shows the results. When evaluated with a linear probe, Seurat regression and CycleVI remove cell cycle information comparably, with macro F1 scores for phase prediction of 0.274 and 0.270, respectively, compared to 0.539 for the uncorrected embedding. Under a nonlinear probe, however, the two methods diverge sharply: phase prediction remains high after Seurat regression (F1 = 0.516, close to the uncorrected value of 0.574), while CycleVI retains far less phase information (F1 = 0.342). To rule out drug-induced confounding of this comparison, we repeated the analysis using only vehicle-treated cells, in which no drug response can contribute to phase structure. The pattern persisted: Seurat regression reduced nonlinear phase predictability only marginally (F1 = 0.598 vs. 0.607 uncorrected), while CycleVI produced a substantially cleaner residual (F1 = 0.425). This confirms that linear regression, by construction, removes only the linear projection of cycle variance onto the cycle scores, while the underlying circular structure, which are invisible to linear probes but recoverable by neighborhood-based analyses such as clustering, UMAP, and trajectory inference, remains largely intact.

**Figure 7:**
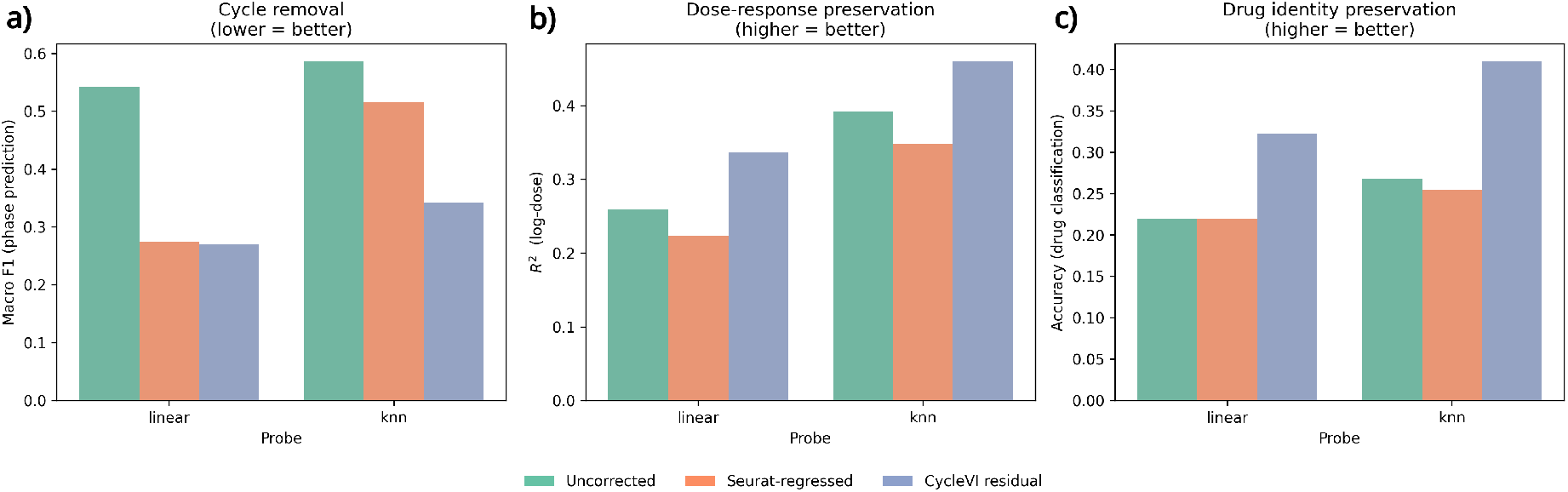
CycleVI disentangles cell cycle from drug response more effectively than linear cycle regression. Three representations of A549 lung adenocarcinoma cells from the sci-Plex3 dataset, treated with 9 HDAC inhibitors across four doses alongside vehicle controls, are compared at matched dimensionality: uncorrected log-normalized expression followed by PCA (green), the same pipeline with prior Seurat regression of S and G2/M cycle scores (orange), and CycleVI’s residual latent space (blue). Cross-validated linear and nonlinear (k-nearest-neighbor, *k* = 30) probes were trained on each representation.**a)** Macro F1 for cell cycle phase classification (lower = more complete cycle removal). Seurat regression and CycleVI perform comparably under the linear probe, but cycle structure largely persists after Seurat regression under the nonlinear probe (F1 = 0.516) and is substantially reduced only by CycleVI (F1 = 0.342). **b)** Cross-validated *R*^2^ for log-dose regression (higher = better dose-response preservation). **c)** Cross-validated accuracy for 9-class drug classification within treated cells (higher = better drug-identity preservation). In both b) and c), CycleVI exceeds the uncorrected baseline, indicating that removing cycle variance actively elucidates drug-response structure rather than merely preserving it.

CycleVI’s more complete cycle removal also improves the recovery of drug-related biology (Figure 7b and c). Cross-validated *R*^2^ for log-dose regression was 0.259 in uncorrected PCA, 0.224 after Seurat regression, and 0.336 in CycleVI’s residual space under the linear probe, with the same ordering under the nonlinear probe (0.387, 0.348, and 0.460 respectively). Classification accuracy for 9 drug identities within treated cells showed the same pattern, reaching 0.322 (linear) and 0.410 (nonlinear) in CycleVI’s residual space compared to 0.220 and 0.255 after Seurat regression. Notably, CycleVI exceeds even the uncorrected baseline on these biology-preservation metrics. This reflects an expected consequence of successful disentanglement: when cycle variance is removed from the representation without attenuating the drug response, the latter occupies a larger fraction of the low-dimensional embedding and becomes more readily accessible to downstream models.

Together, these results demonstrate that linear cycle regression and CycleVI differ qualitatively, not just quantitatively. Seurat regression decouples phase inference from cycle correction: a fixed marker-based phase estimate is computed once and then linearly subtracted from each gene. CycleVI instead solves both problems jointly, and this joint optimization allows the phase estimate to be informed by the data the correction will be applied to and forces cycle-related variance into the dedicated circular subspace by construction. As a consequence, CycleVI removes more of the circular cycle geometry and clarifies drug-response signals that are partially obscured by proliferation in the uncorrected data.

### 3.6 CycleVI spatially resolves proliferative heterogeneity in a metastatic breast cancer biopsy

Understanding the spatial organization of cellular proliferation within the tumor microenvironment is a central goal in cancer biology [36], yet it remains a significant technical challenge. Single-cell methods based on scRNA-seq can profile proliferative states but lose the native architectural context. Spatial transcriptomics platforms recover this context but introduce technical constraints that complicate cell-cycle inference. Slide-seq, in particular, produces sparse profiles with low transcript counts per bead [37], and does not always achieve single-cell resolution, with each bead potentially containing transcripts from more than one cell. These limitations make Slide-seq a demanding test case for cell-cycle inference. CycleVI is a useful candidate here because its variational framework explicitly models count-level noise and its disentanglement architecture absorbs non-cycle heterogeneity into a separate latent space. Here we apply CycleVI to a Slide-seq dataset from a metastatic breast cancer biopsy from Klughammer et al. [38] to determine if it could infer and map cell cycle states directly within the tissue architecture.

The analysis focuses on a liver metastasis biopsy containing a well-defined tumor region surrounded by non-tumor liver parenchyma, as confirmed by Hematoxylin and Eosin (H&E) staining of a serial tissue section (Figure 8a). CycleVI was run on the bead-level expression matrix, which contains transcriptomic information from thousands of discrete 10*µm* locations across the tissue slice, to infer a continuous cell cycle phase angle for each bead. This inference was performed without any prior information regarding the histological identity or location of the beads.

**Figure 8:**
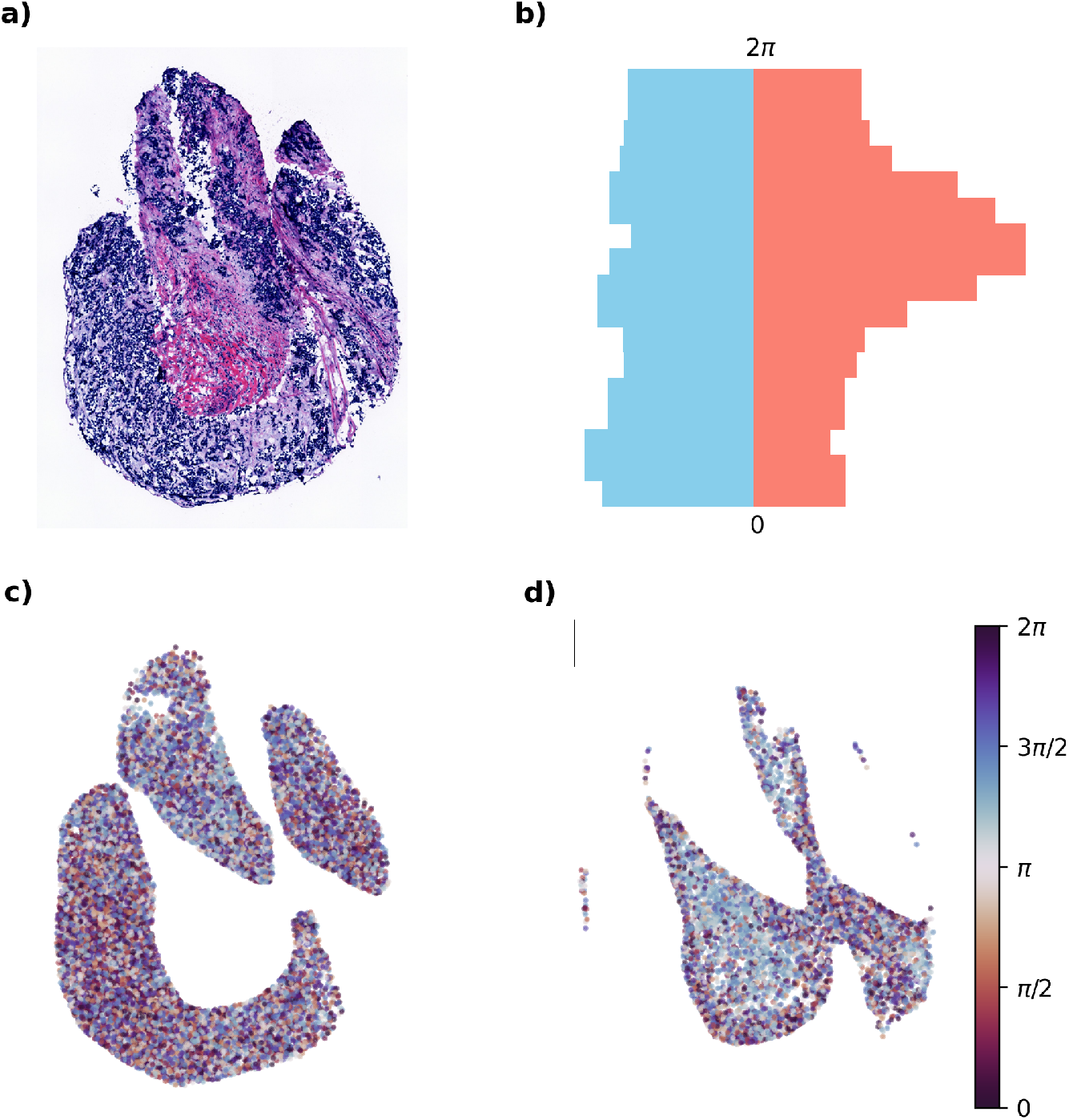
CycleVI spatially resolves proliferative heterogeneity in a metastatic breast cancer biopsy. **a)** Hematoxylin and Eosin (H&E) stain of a serial section from the metastatic breast cancer biopsy (liver metastasis), showing the histologically distinct tumor region (dark purple) and the surrounding non-tumor liver parenchyma. **b)** Distribution of CycleVI-inferred cell cycle phase angles for all Slide-seq beads, plotted as a back-to-back histogram. The vertical axis is the cell cycle angle, ranging from 0 to 2*π*. The horizontal axis is bead density, with tumor beads (blue) extending to the left of the central axis and non-tumor beads (red) extending to the right. Beads within the tumor exhibit a broad distribution across all phases, indicative of active proliferation, while non-tumor beads are strongly concentrated in the G1 phase (histogram peak), characteristic of quiescence. **c)**-**d)** Spatial map of the Slide-seq beads, with each bead colored by its CycleVI-inferred phase angle according to the provided color key. The two panels show tumor (c) and non-tumor (d). The tumor region displays a salt-and-pepper pattern of all colors, confirming the co-localization of cells in diverse cycle stages. In contrast, the non-tumor region is evenly colored, corresponding to the G1 phase. We see CycleVI’s ability to map dysregulated proliferation in its native spatial context using static transcriptomic data.

Mapping the inferred phase angle of each bead back to its spatial coordinates revealed a striking pattern of proliferative heterogeneity that precisely mirrored the underlying histology (Figure 8c-d). The region corresponding to the tumor mass appeared as a mosaic of colors, representing the full spectrum of cell cycle phases. This spatial pattern indicates that tumor cells are asynchronously cycling, with cells at all stages of division co-existing in close proximity. In stark contrast, beads located in the nontumor region, corresponding to the liver parenchyma, were primarily in the range associated with the G1 phase.

This visual observation was quantified by plotting the distribution of inferred phase angles for beads stratified by their histological location (Figure 8b). The distribution for beads within the tumor region is broad, with substantial populations present across all phases of the cell cycle, confirming high and continuous proliferative activity. Conversely, the distribution for beads in the non-tumor region is sharply peaked in G1, a profile characteristic of a largely quiescent or terminally differentiated cell population.

This result provides a clear, spatially resolved confirmation of a fundamental hallmark of cancer: dysregulated and uncontrolled cell proliferation. The analysis demonstrates that CycleVI can, from transcriptomic data alone, quantitatively distinguish the highly proliferative niche of the tumor from the quiescent state of the surrounding tissue. The model’s ability to recover this core biological principle in situ serves as a strong orthogonal validation of its biological relevance and accuracy.

Furthermore, the success of this application highlights the robustness of the CycleVI framework. Slide-seq data is known to be sparser than typical scRNA-seq, with fewer unique transcripts detected per observation (bead). The ability of CycleVI’s model to infer a coherent and biologically meaningful cell cycle trajectory from this sparser data underscores its resilience to technical noise and its applicability to challenging, real-world datasets. This analysis showcases CycleVI’s capability to bridge molecular-level inferences with tissue-level architecture, facilitating the study of the spatial dynamics of cell proliferation in complex tissues.

### 3.7 CycleVI disentanglement clarifies the human hematopoietic differentiation manifold

Having established in Section 3.5 that CycleVI removes cell-cycle variation more completely than linear regression, we now ask whether this improved disentanglement has measurable downstream consequences for biological analyses where the cell cycle is a confounder rather than the signal of interest. Human hematopoiesis, the process by which hematopoietic stem cells (HSCs) differentiate into all mature blood lineages, serves as a perfect example of such a system. In the bone marrow, progenitor cells must proliferate to expand and generate progeny, while simultaneously undergoing transcriptional changes that commit them to specific cell fates. The strong, often dominant, transcriptional signature of the cell cycle can therefore obscure the more subtle signals that define differentiation trajectories, leading to erroneous interpretations of lineage relationships.

To evaluate CycleVI’s performance in this challenging context, we applied it to a public single-cell RNA-seq dataset of 31,634 human bone marrow cells enriched for the CD34+ hematopoietic stem and progenitor cell (HSPC) population [39]. This dataset was specifically generated to capture the continuous landscape of early hematopoietic differentiation and is known to have a strong cell cycle signal, and the original authors noted the necessity of a correction step, stating that “expression of cell cycle genes can confound the ordering of cells in a differentiation trajectory.”

As shown in Figure 9, this confounding effect is apparent. A UMAP projection of the raw, log-normalized expression data reveals that cells are organized strongly by their position in the cell cycle (Supplementary Figure S3, top), causing cells from distinct lineages to cluster together and obscuring the underlying differentiation manifold. Applying a standard correction method, which involves regressing out cell cycle scores as implemented in Seurat, only partially mitigates this issue and fails to resolve the distinct progenitor populations clearly.

**Figure 9:**
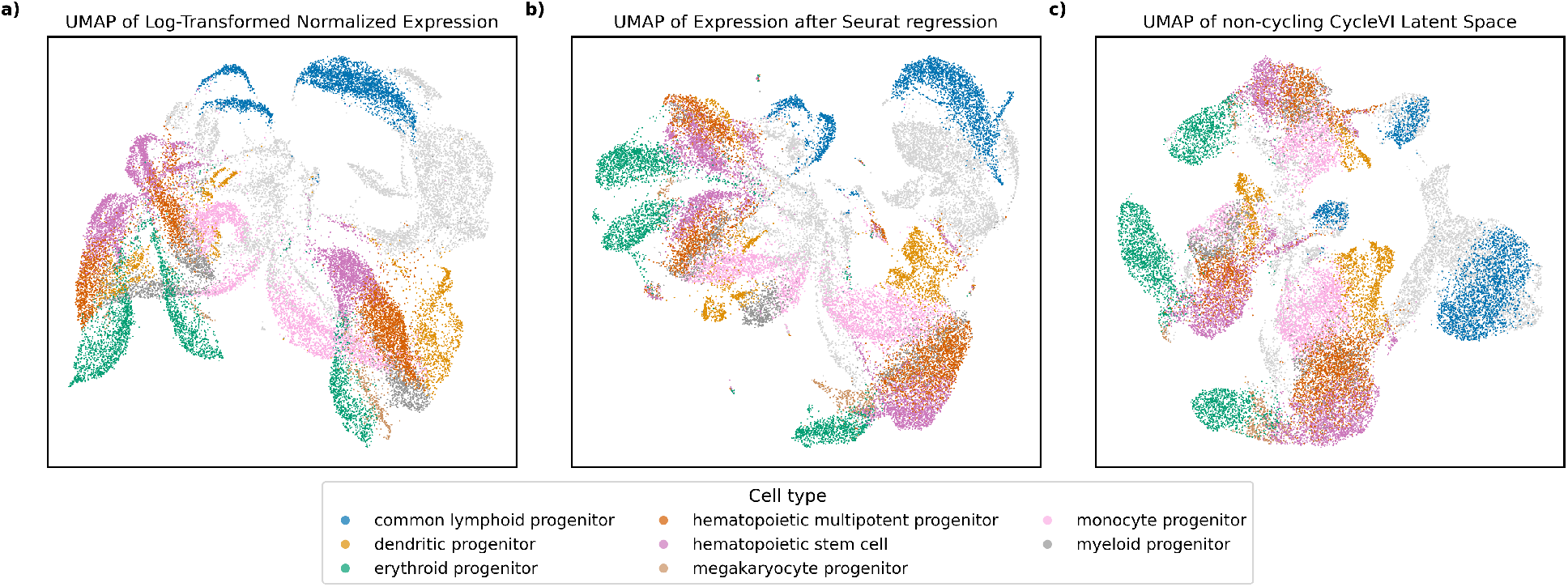
Principled disentanglement by CycleVI clarifies the human hematopoietic differentiation manifold. Analysis of human hematopoietic stem and progenitor cell (HSPC) differentiation is a canonical challenge where the strong transcriptional signature of proliferation masks the subtle signals of lineage commitment. We applied CycleVI to a dataset of 31,634 human bone marrow HSPCs from three distinct donors, a system where the original authors noted the necessity of cell cycle correction. **a)** A UMAP projection of the raw log-normalized expression data shows that cells are organized primarily by cell cycle phase, confounding the underlying differentiation structure. **b)** Applying a standard correction method by regressing out cell cycle scores (Seurat) provides only a partial improvement, with lineage trajectories remaining poorly resolved. **c)** In contrast, the UMAP of CycleVI’s residual latent space reveals a well-structured differentiation landscape. The model’s principled isolation of cell cycle effects clearly resolves trajectories from multipotent progenitors toward committed erythroid, myeloid, and lymphoid fates. This disentangled representation also unmasks significant inter-donor variation as a major source of heterogeneity, demonstrating the model’s ability to enhance biologically relevant signals beyond the primary confounder.

In contrast, the UMAP based on CycleVI’s residual latent space, the representation from which cell cycle variation has been disentangled, reveals a well-structured differentiation landscape. The structure of the disentangled embedding recapitulates the canonical organization of the hematopoietic hierarchy. The most prominent feature of panel (c) is the separation of cells into three disconnected regions, which correspond to the three donors in the dataset (Supplementary Figure S3, bottom). Within each donor, hematopoietic stem cells (light purple) and multipotent progenitors (brown) occupy a region from which three lineage trajectories extend: a lymphoid branch terminating in the common lymphoid progenitors (blue), an erythroid branch with the associated megakaryocyte progenitors (green and tan), and a myeloid branch comprising myeloid (grey), monocyte (light pink), and dendritic (yellow) progenitors. This is the canonical structure described by the original authors of the dataset, which the cell-cycle signal had obscured in panels (a) and (b). One detail of this representation is worth highlighting. Lymphoid progenitors in this dataset are biased toward G1/G0 compared to the other lineages (Supplementary Figure S3, top), so their identity correlates with cell-cycle phase. CycleVI nonetheless resolves them as a distinct cluster in the residual space because the adversarial classifier is annealed to zero over training, allowing biologically meaningful cell-type structure to be retained even when it correlates with cycle state.

By isolating the cell cycle signal into its dedicated latent subspace, CycleVI preserves the integrity of the differentiation-related transcriptional programs. This demonstrates the significant advantage of a principled disentanglement approach over regression-based methods, enabling a clearer and more accurate analysis of complex, concurrent biological processes.

## 4 Conclusion

In this work, we introduce CycleVI, an interpretable deep generative model for disentangling cell cycle–driven expression from other sources of variation in single-cell transcriptomics. Cell cycle progression often constitutes the dominant transcriptional signal, masking subtler variation related to cell identity, differentiation, or perturbation response and complicating downstream analyses. CycleVI represents a conceptual shift from confounder correction toward principled isolation of proliferative variation and further distinguishes itself from other models by neither discretizing the cell cycle nor relying on RNA velocity.

A key strength of CycleVI lies in its foundation on variational inference, which is well-suited for the inherent stochasticity of gene expression. Even cells occupying the same cell cycle phase can exhibit substantial variability in phase marker expression, and the probabilistic framework of variational autoencoders enables robust modeling of this noise in high-dimensional data. Consequently, CycleVI recovers a representation of cell cycle progression that is resilient to transcriptional noise, distinguishing it from more deterministic or heuristic approaches.

The success of CycleVI stems from its biologically informed architecture designed for principled disentanglement. Its key innovations include a partitioned latent space that explicitly dedicates a circular subspace to the cell cycle, a dual-decoder system that uses a gene-specific Fourier series to model periodic expression, and an adversarial classifier that enforces a clean separation of signals.

The model accurately recovers known cell cycle transcriptional dynamics, including phase-specific expression of canonical markers, histone accumulation during S phase, ribosomal protein dynamics, and post-mitotic transcript clearance. Benchmarking against an orthogonal FUCCI protein-level readout showed that CycleVI outperformed seven established cell cycle inference methods on both a controlled pulse subset and a more heterogeneous full dataset. Importantly, CycleVI retained performance when experimental heterogeneity increased, whereas several competing methods degraded substantially.

CycleVI also distinguished cycling from non-cycling populations in complex tissue settings. In FFPE breast cancer tissue, immune populations were concentrated in the G1/G0-associated region of the inferred cycle, whereas tumor populations were distributed broadly across cell cycle phases, consistent with asynchronous proliferation. In spatial transcriptomics data from a metastatic breast cancer biopsy, CycleVI mapped proliferative heterogeneity directly onto tissue architecture, separating broadly cycling tumor regions from largely quiescent surrounding tissue.

A central aim of CycleVI is not only to infer cell cycle phase, but to remove its confounding influence from representations used for downstream analysis. In the sci-Plex3 drug perturbation benchmark, CycleVI removed nonlinear cell cycle structure more effectively than standard Seurat-style linear regression of S and G2/M scores. At the same time, CycleVI preserved and clarified drug-response information, improving recovery of both dose and drug identity relative to uncorrected and linearly regressed representations. Finally, CycleVI clarified the differentiation manifold of human hematopoietic stem and progenitor cells, a setting in which proliferation and lineage commitment are strongly intertwined. This illustrates the utility of CycleVI for systems in which the cell cycle is neither purely nuisance variation nor the primary biological question, but a confounding process that overlaps with other transcriptional programs.

A comparison with the more generic scVI model highlights a crucial trade-off between flexibility and generalization. While scVI achieves lower reconstruction error on training data, CycleVI performs better on unseen validation and test sets, indicating that its architectural constraints act as an effective form of regularization. This supports a broader principle in computational biology: incorporating domain-specific inductive biases into deep learning models can improve generalization and reduce overfitting, in addition to enhancing interpretability.

In summary, CycleVI provides a robust and interpretable framework for separating proliferative transcriptional programs from other sources of variation in static scRNA-seq data. While limitations remain, including reliance on heuristic initialization, the lack of explicit modeling of quiescent states, and its reliance on larger datasets, CycleVI’s ability to clarify cellular heterogeneity and its improved generalization highlight the value of integrating domain knowledge into deep learning frameworks. We anticipate that these ideas will extend to future generative models aimed at deconstructing the complex, overlapping transcriptional programs that orchestrate cellular identity.

## Code availability

CycleVI is available at www.github.com/jeuken/CycleVI, and archived under the DOI doi.org/10.5281/zenodo.19919729.

## Data availability

All datasets analyzed here are publicly available through their original publications. The human fetal lung fibroblast (GSE167609), FUCCI-labeled cell (GSE128365), FFPE breast cancer tissue (GSE243280), and sci-Plex3 (GSM7056150) datasets are available from the GEO database. The human hematopoietic progenitor dataset is available from the Single Cell Expression Atlas under accession E-HCAD-6. The metastatic breast cancer spatial transcriptomics dataset is available from the Single Cell Portal under accession SCP2702.

