## Supplementary information for "CycleVI: Isolating cell cycle variation with an interpretable deep generative model"

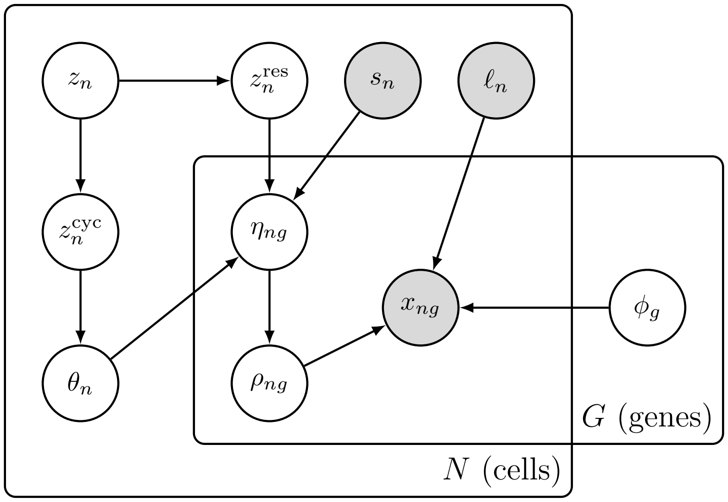


Figure S1: **A plate diagram of the CycleVI generative model.** A probabilistic graphical model representation (plate diagram) of the generative process for CycleVI, as described in detail in Section 2.1. The model maps the gene expression matrix ($x$) to a partitioned latent space ($z$) and learns the parameters of a generative distribution $p(x|z)$

.

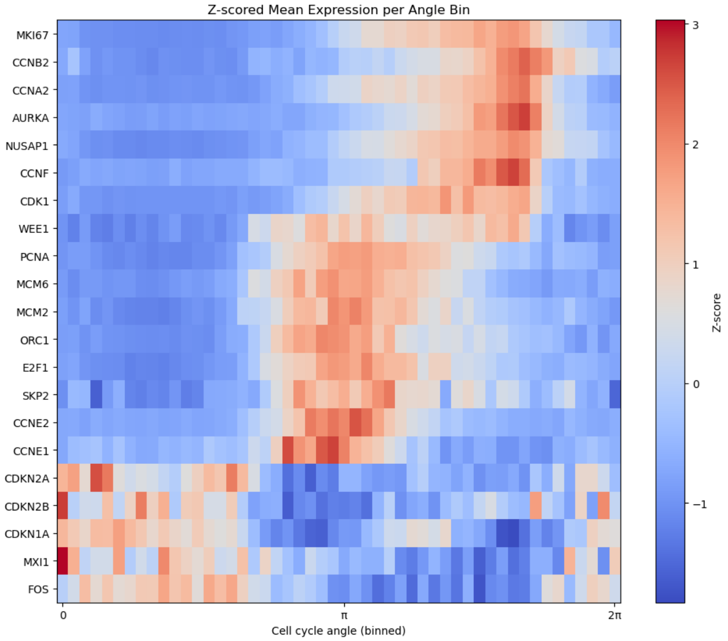


Figure S2: **CycleVI-inferred phase angle correctly organizes the temporal expression cascade of canonical cell cycle marker genes.** Heatmap displaying the mean, Z-scored expression of a panel of canonical cell cycle marker genes, plotted against the continuous cell cycle phase angle inferred by CycleVI. Gene expression data is from the human fetal lung fibroblast dataset. Each row represents a single gene, and rows are manually ordered to highlight the known temporal progression of expression. The x-axis represents the inferred cell cycle phase, which has been binned for visualization. The color scale represents the Z-scored mean expression within each angle bin.

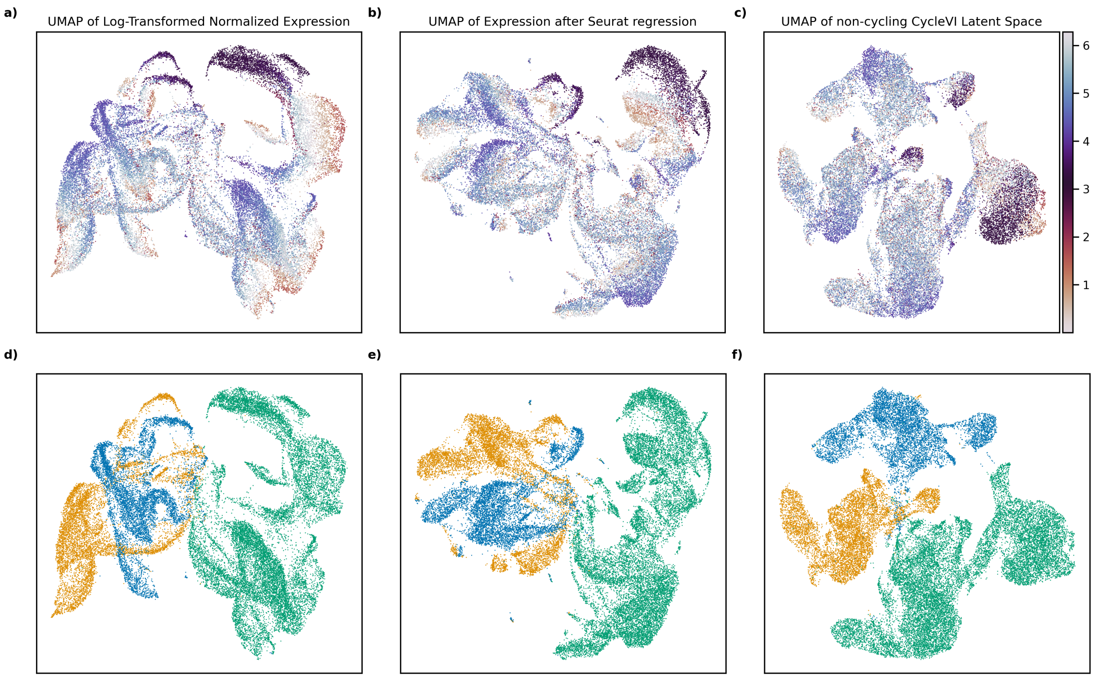


Figure S3: **CycleVI successfully disentangles cell cycle variation from complex inter-donor variation in hematopoietic progenitors.** Comparison of UMAP projections of the human hematopoietic progenitor dataset (Setty et al. ), demonstrating the success of different methods to remove the confounding cell cycle signal while preserving other biological variation. This figure provides supporting evidence for the analysis in main text Section 3.6. **Top row (panels a-c)**: UMAPs colored by the CycleVI-inferred cell cycle phase angle. A continuous, circular color map (from 0 to $2\pi$) indicates the presence of a cell cycle signal. **Bottom row (panels d-f)**: The exact same UMAP projections as the top row, but colored by the sample donor (three distinct donors), illustrating the presence of inter-donor variation. **Panels (a) and (d)**: UMAP of the raw, log-normalized gene expression. The visualization is dominated by the cell cycle (a), which forms a distinct circular structure that confounds and obscures the underlying inter-donor variation (d). **Panels (b) and (e)**: UMAP of the data after applying a standard cell cycle correction workflow (Seurat-based regression of S and G2/M scores), as described in the main text. The cell cycle signal is reduced but still clearly present, as seen by the non-uniform grouping of phase-specific cells (b). This remaining artifact continues to confound the inter-donor variation, which remains poorly resolved (e). **Panels (c) and (f)**: UMAP of the 8-dimensional CycleVI residual latent space ($z^{res}$). This representation is now demonstrably free of the cell cycle signal, as shown by the uniform mixing of phase angles (c). As a result, the inter-donor variation, which was previously masked, now emerges as a key axis of separation (f).

### 1 Datasets

To comprehensively evaluate the performance, robustness, and utility of CycleVI, we selected a diverse set of six publicly available single-cell RNA-seq datasets. These datasets span different biological systems, technical platforms, and analytical challenges, allowing us to test specific capabilities of the model, from the accuracy of phase inference to the biological relevance of the disentangled latent space.

**Human fetal lung fibroblasts**: To benchmark the accuracy of CycleVI's phase inference against state-of-the-art methods, we used a dataset of 5,367 human fetal lung fibroblast cells, previously analyzed in the DeepCycle study [1]. This dataset is well-suited for comparative analysis as it contains a large, actively proliferating cell population for which a continuous cell cycle phase has been previously inferred using a velocity-based method, providing a strong reference for comparison.

**Human hematopoietic progenitors**: To test CycleVI's ability to disentangle the cell cycle from a complex, concurrent biological process, we used the human hematopoietic dataset from Setty et al. [2]. This dataset consists of 31,634 bone marrow cells from healthy human donors, enriched for CD34+ hematopoietic stem and progenitor cells (HSPCs). It captures a continuous differentiation landscape from multipotent progenitors to committed lymphoid, erythroid, and myeloid lineages. As proliferation is tightly coupled with differentiation in this system, it represents a canonical "stress test" for any method aiming to isolate cell cycle effects without distorting the underlying differentiation trajectories.

**HDAC inhibitor chemical perturbation screen (sci-Plex3)**: To benchmark CycleVI against linear cell cycle regression in a setting where the biological signal of interest is partially entangled with the cell cycle, we used a subset of the sci-Plex3 chemical transcriptomics dataset from Srivatsan *et al* [3]. We selected A549 lung adenocarcinoma cells treated with 9 HDAC inhibitors (Pracinostat, Abexinostat, Belinostat, Mocetinostat, Dacinostat, Givinostat, Tucidinostat, CUDC-907, Quisinostat) at four doses (10 nM, 100 nM, 1 $\mu$M, and 10 $\mu$M) alongside DMSO vehicle controls. HDAC inhibitors induce G1 and G2/M cell cycle arrest and modulate the expression of canonical cycle genes such as CDKN1A and E2F targets, so their transcriptional response shares structure with the cell cycle program. This makes the dataset particularly suited for evaluating whether a cycle correction method preserves or distorts a biological signal of interest.

**FUCCI-labeled cells**: To validate the inferred cell cycle phase against a non-transcriptional, orthogonal ground truth, we used the dataset from Battich et al. [4]. This study employed the Fluorescent Ubiquitination-based Cell Cycle Indicator (FUCCI) system, which uses the fluorescence of two post-translationally regulated proteins to provide a continuous, quantitative measure of each cell's position in the cycle. This protein-level measurement provides an unbiased benchmark to assess the accuracy of phase inference from transcriptomic data alone.

**FFPE breast cancer tissue**: To assess the model's robustness on technically challenging samples and its ability to distinguish cycling from non-cycling populations, we used a dataset of a Formalin-Fixed Paraffin-Embedded (FFPE) breast cancer biopsy from Janesick et al. [5]. This dataset contains a mix of mitotically active tumor cell populations and quiescent immune cell infiltrates, providing a real-world test case for the model's performance in a complex tissue microenvironment.

**Metastatic breast cancer spatial transcriptomics**: To demonstrate CycleVI's utility on spatially resolved transcriptomics data and in a clinically relevant context, a publicly available Slide-seq dataset from a metastatic breast cancer (MBC) core needle biopsy was analyzed. The data originates from the Klughammer et al. [6] multi-modal atlas of MBC, specifically from a liver metastasis of an HR+/HER2- tumor. Slide-seq provides whole-transcriptome data with near-cellular resolution using $10\mu m$ barcoded beads, with expression measured as UMI counts. The dataset consists of gene expression counts for thousands of beads, each with associated spatial coordinates. For this analysis, the provided Hematoxylin and Eosin (H&E) stain of a serial tissue section was used to manually delineate the primary tumor region from the surrounding non-tumor liver parenchyma, allowing for the stratification of the Slide-seq beads by their histological context.

### 2 Selection of the number of harmonics $K$

To select the number of harmonics K in the Fourier decoder, we trained CycleVI with varying $K \in 1, ..., 5$ on the hematopoiesis dataset and transferred only the fitted gene-specific Fourier coefficients to the FUCCI dataset. The encoder, residual decoder, and latent representations were re-fit on FUCCI, so that the Spearman correlation between the inferred phase and the FUCCI-derived ground truth measures exclusively whether the learned periodic expression profiles generalize across datasets. Coefficients that capture genuine cell-cycle biology should transfer, those that overfit dataset-specific noise should not. Performance increased from $K = 1$ ($\rho = 0.316$) through $K = 3$ ($\rho = 0.393$), plateaued at $K = 4$ ($\rho = 0.403$), and dropped sharply at $K = 5$ ($\rho = 0.342$). The plateau at $K = 3$ to $K = 4$ indicates that three or four harmonics are sufficient to capture the periodic structure shared across cell types, while the collapse at $K = 5$ confirms that additional harmonics fit dataset-specific noise that does not transfer. We selected K=3 as the most parsimonious choice within the plateau region, with 6 Fourier coefficients per gene compared to 8 for $K = 4$, and sitting further from the overfitting regime indicated by $K = 5$.

### 3 CycleVI produces stable phase inferences that are robust to noisy initialization

We assessed the robustness of our method through two tests on a dataset of 31,634 cycling cells from Setty et al. [2]. First, to evaluate the stability of the inference process, we executed the model five independent times on the same dataset. The inferred cell cycle phases showed great consistency across runs. Even the worst-performing pairwise comparison yielded a Spearman correlation of 0.937, indicating a highly stable solution (Figure S4a). Second, we tested the model's resilience to the quality of the initial phase estimates by introducing substantial random noise, equivalent to 30% of the full $[0, 2\pi]$ range, to the initialization angles. Despite this significant perturbation, the final inferred phases maintained a high degree of agreement with the original results, achieving a Spearman correlation of 0.912 (Figure S4b). Together, these results demonstrate that our method is highly robust, producing reproducible phase inferences that are insensitive to both stochastic run-to-run variability and noisy initialization.

### 4 CycleVI phase inference aligns with velocity-based methods using only static expression data

For each cell, CycleVI infers a phase angle between 0 and $2\pi$, representing its position along the cell cycle continuum. To evaluate these estimates, we compared the phases inferred by CycleVI, Seurat, and DeepCycle using the same human fetal lung fibroblast dataset originally used in the DeepCycle study [1]. Figure S5 contrasts CycleVI's output with two references: (1) the phase angle computed from Seurat's G2/M and S scores that was quantile-transformed to follow a uniform distribution over $[0, 2\pi]$, which is used as a weak prior during CycleVI training (see Section 2.1.3), and (2) the phase predicted by DeepCycle. For easier comparison, DeepCycle's output in $[0, 1]$ was scaled to $[0, 2\pi]$ and circularly shifted by 2.9 radians to align trajectories across models.

The Seurat-derived phase closely aligns with the CycleVI-inferred one. This can be expected, as CycleVI is initialized using the Seurat results. However, the model retains flexibility to deviate from this initialization, showing that the weak prior does not rigidly constrain learning and allows CycleVI to discover its own optimal ordering. This is evident by the fact that the inferred phase shows a much stronger agreement with DeepCycle, suggesting that it is possible to recover meaningful cell cycle trajectories without relying on RNA velocity information derived from the ratio of spliced to unspliced RNA. Instead, static gene expression snapshots alone appear sufficient to capture the progression through the cell cycle. It is important to note that all three methods infer pseudotime trajectories. These represent each cell's relative position along the cell cycle, rather than actual time. As a result, comparisons between methods

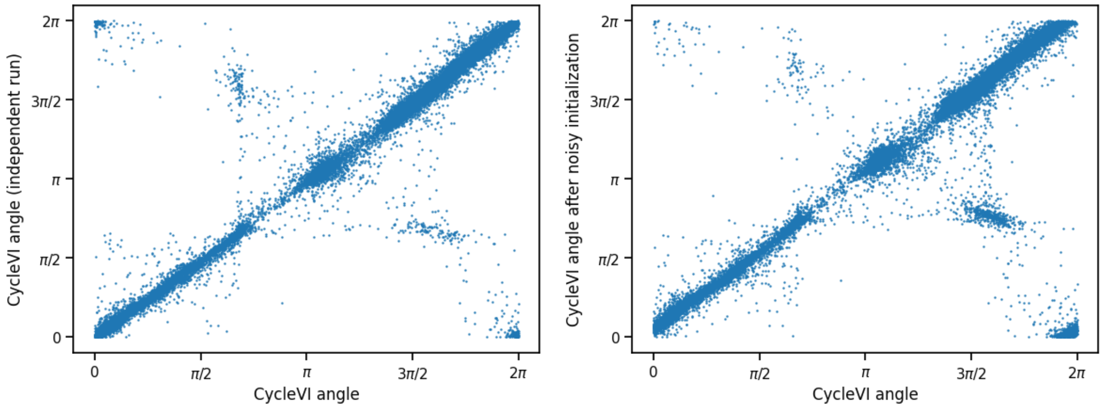


Figure S4: **CycleVI demonstrates high robustness to run-to-run variability and initialization noise.** Robustness analysis of the CycleVI model performed on the human hematopoietic progenitor dataset (Setty et al., 31,634 cells). Both plots are scatterplots comparing inferred cell cycle angles (in radians) from two different model runs. **Left**: Assessment of stochastic run-to-run variability. The plot shows the pairwise comparison of inferred angles from two of five independent model runs. This pair was selected as it represents the lowest concordance (i.e., the worst-performing pair) out of 20 pairwise comparisons. The high Spearman correlation (Spearman's $\rho = 0.937$) demonstrates that the model optimization is highly stable and consistently converges to the same solution, overcoming the challenges of non-convexity in the optimization landscape. **Right**: Assessment of sensitivity to initialization noise. The plot compares the inferred angles from a standard model run (x-axis) against a run where the Seurat-derived initialization angles ($\theta_n^{ref}$) were perturbed with significant uniform white noise (magnitude of $0.3 \times 2\pi$, or 30% of the full range). The resulting high correlation (Spearman's $\rho = 0.912$) demonstrates that the model is robust to noisy priors. This confirms that the final phase inference is not rigidly determined by the initialization, but is rather learned from the underlying gene expression data.

are expected to show monotonic, but potentially nonlinear, relationships.

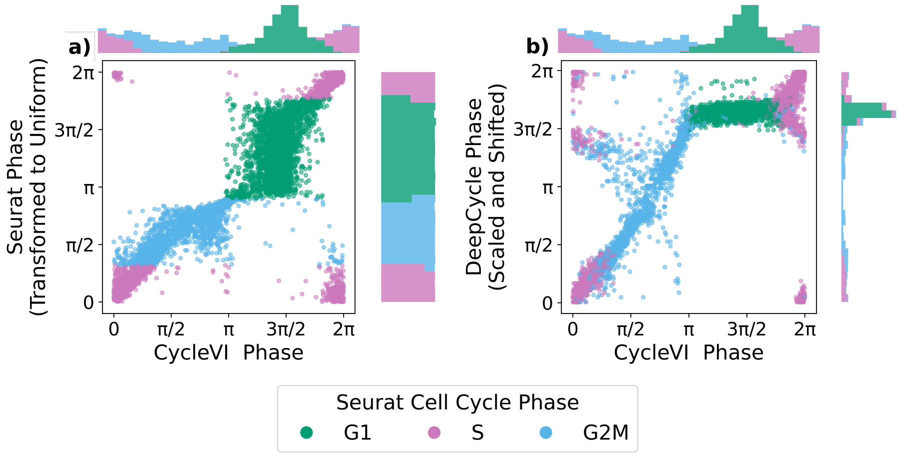


Figure S5: **Comparison of the cell cycle phase inferred by CycleVI, Seurat, and DeepCycle. a)** Scatterplot of the quantile-transformed Seurat phase derived from G2/M and S scores, mapped to a uniform distribution over $[0, 2\pi]$, versus CycleVI-inferred phase. **b)** Scatterplot of the DeepCycle phase, scaled to $[0, 2\pi]$ and circularly shifted by 2.9 radians, versus CycleVI-inferred phase. Marginal histograms show the phase distributions colored by Seurat-assigned cell cycle phase labels.

### 5 Biologically informed architecture enhances model generalization

Finally, we assessed the technical benefits of incorporating a strong, biologically informed inductive bias into our model. An important question is whether constraining the VAE with a specialized cell cycle decoder might impair its overall ability to reconstruct the transcriptome compared to a more flexible, generic model. To address this, we compared CycleVI directly against scVI, an equivalent VAE that does not include this specialized decoder.

We trained both models on the hematopoiesis data from Setty et al. [2] for 400 epochs with 10 latent dimensions. We did so 20 times, and in each run we train the model in a random sample of 10% of the data, using another independent sample of 10% as the test set. As shown in Figure S8, the results reveal a classic and important trade-off between model flexibility and generalization. While scVI's unconstrained architecture allowed it to achieve a lower reconstruction loss on the training sets, CycleVI shows a superior performance on the unseen test sets.

This result is significant. It indicates that the greater flexibility of the stan-

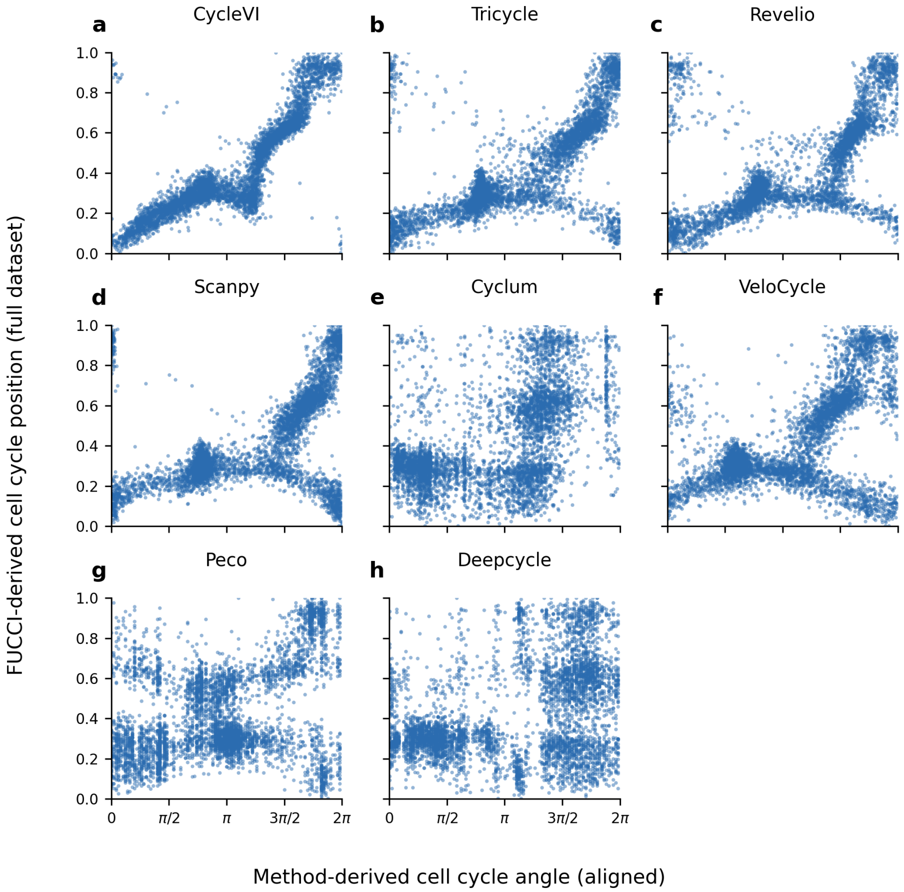


Figure S6: **Method-derived cell cycle angle versus FUCCI-derived cell cycle position across the full RPE1 benchmark dataset.** Each panel (a–h) shows one method's predicted cell cycle angle (x-axis, aligned to FUCCI by optimal circular shift and direction) plotted against the FUCCI-derived reference position (y-axis) for all 4,955 cells.

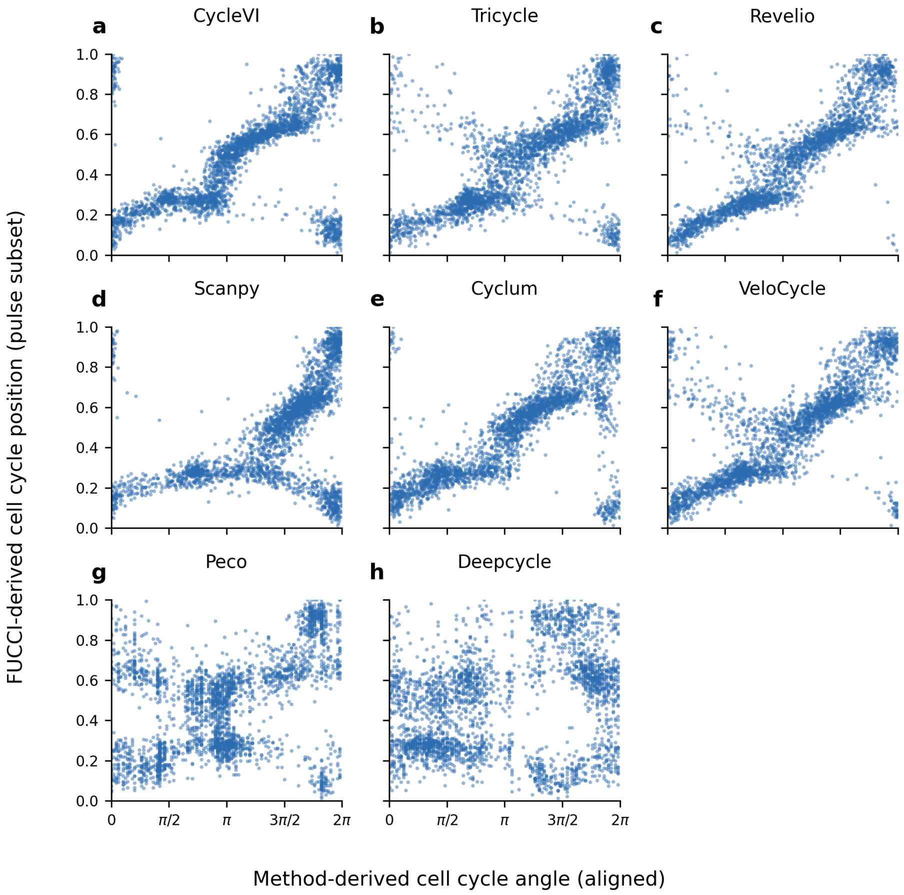


Figure S7: **Method-derived cell cycle angle versus FUCCI-derived cell cycle position on the pulse subset of the RPE1 benchmark dataset.** As in Supplementary Figure S6, but restricted to pulse-condition cells.

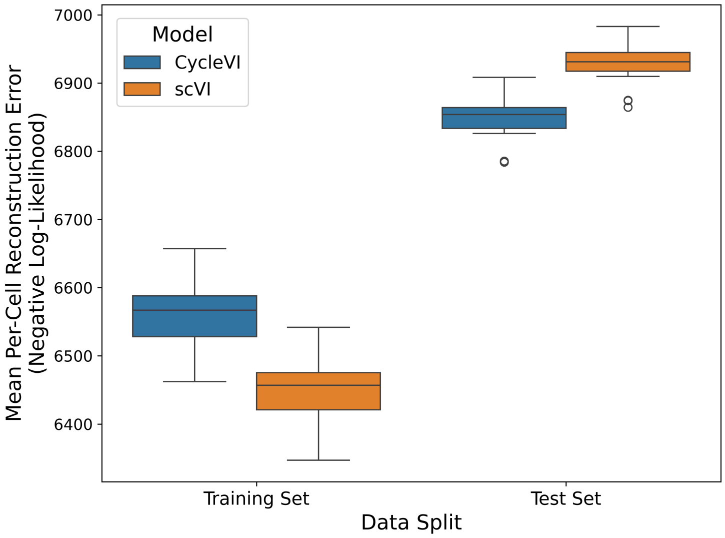


Figure S8: **Reconstruction performance of scVI and CycleVI.** Boxplots showing the mean per-cell reconstruction error for scVI and CycleVI on the hematopoiesis dataset from Setty et al. Each box summarizes results across 20 runs with different random data samples (10% for training, 10% for testing). Both models were trained with 10 latent dimensions for 400 epochs. The reconstruction error is defined as the negative log-likelihood of observed gene expression under the predicted negative binomial distribution, summed over all genes and averaged across cells.

dard VAE architecture leads to minor overfitting on the training data. In contrast, the architectural constraints of CycleVI act as an effective regularizer, improving the model's ability to generalize to new data. This provides evidence that incorporating explicit biological knowledge does not compromise the model's reconstructive power, but rather it provides a powerful inductive bias that guides the model toward learning more robust and generalizable representations of cellular transcription programs.

### 6 Implementation

The model was implemented using the scvi-tools library [7]. Data preprocessing and analysis, including Seurat's cell cycle scoring, were performed using scanpy [8]. The cell cycle genes used in such scoring are the ones provided by Seurat, taken from Tirosh et al. [9]. When necessary, these gene names were converted into Ensembl IDs using BioMart [10].

[6] Johanna Klughammer, Daniel L Abravanel, Åsa Segerstolpe, Timothy R Blosser, Yury Goltsev, Yi Cui, Daniel R Goodwin, Anubhav Sinha, Orr

Ashenberg, Michal Slyper, et al. A multi-modal single-cell and spatial expression map of metastatic breast cancer biopsies across clinicopathological features. *Nature Medicine*, 30(11):3236–3249, 2024.

[7] Adam Gayoso, Romain Lopez, Galen Xing, Pierre Boyeau, Valeh Valiollah Pour Amiri, Justin Hong, Katherine Wu, Michael Jayasuriya, Edouard Mehlman, Maxime Langevin, Yining Liu, Jules Samaran, Gabriel Misrachi, Achille Nazaret, Oscar Clivio, Chenling Xu, Tal Ashuach, Mariano Gabitto, Mohammad Lotfollahi, Valentine Svensson, Eduardo da Veiga Beltrame, Vitalii Kleshchevnikov, Carlos Talavera-López, Lior Pachter, Fabian J. Theis, Aaron Streets, Michael I. Jordan, Jeffrey Regier, and Nir Yosef. A Python library for probabilistic analysis of single-cell omics data. *Nature Biotechnology*, 40(2):163–166, February 2022. Publisher: Nature Publishing Group.

[8] F Alexander Wolf, Philipp Angerer, and Fabian J Theis. Scanpy: large-scale single-cell gene expression data analysis. *Genome biology*, 19(1):15, 2018.

[9] Itay Tirosh, Benjamin Izar, Sanjay M. Prakadan, Marc H. Wadsworth, Daniel Treacy, John J. Trombetta, Asaf Rotem, Christopher Rodman, Christine Lian, George Murphy, Mohammad Fallahi-Sichani, Ken Dutton-Regester, Jia-Ren Lin, Ofir Cohen, Parin Shah, Diana Lu, Alex S. Genshaft, Travis K. Hughes, Carly G. K. Ziegler, Samuel W. Kazer, Aleth Gaillard, Kellie E. Kolb, Alexandra-Chloé Villani, Cory M. Johannessen, Aleksandr Y. Andreev, Eliezer M. Van Allen, Monica Bertagnolli, Peter K. Sorger, Ryan J. Sullivan, Keith T. Flaherty, Dennie T. Frederick, Judit Jané-Valbuena, Charles H. Yoon, Orit Rozenblatt-Rosen, Alex K. Shalek, Aviv Regev, and Levi A. Garraway. Dissecting the multicellular ecosystem of metastatic melanoma by single-cell RNA-seq. *Science*, 352(6282):189–196, April 2016. Publisher: American Association for the Advancement of Science.

[10] Damian Smedley, Syed Haider, Benoit Ballester, Richard Holland, Darin London, Gudmundur Thorisson, and Arek Kasprzyk. Biomart–biological queries made easy. *BMC genomics*, 10(1):22, 2009.
